# Improving inference in environmental surveillance by modeling the statistical features of digital PCR

**DOI:** 10.1101/2024.10.14.618307

**Authors:** Adrian Lison, Timothy R. Julian, Tanja Stadler

**Affiliations:** ETH Zurich, Department of Biosystems Science and Engineering, Zurich, Switzerland; SIB Swiss Institute of Bioinformatics, Lausanne, Switzerland; Eawag, Swiss Federal Institute of Aquatic Science and Technology, 8600 Dubendorf, Switzerland; Swiss Tropical and Public Health Institute, 4123 Allschwil, Switzerland; University of Basel, 4055 Basel, Switzerland

**Keywords:** eDNA, wastewater-based epidemiology, dPCR, limit of detection, limit of quantification

## Abstract

Digital polymerase chain reaction (dPCR) is a powerful technique for quantifying gene targets in environmental samples, with various applications such as biodiversity monitoring and wastewater-based epidemiology. However, statistical analyses of environmental dPCR data often assume, explicitly or implicitly, that concentration measurements have a normal or log-normal error structure, which does not reflect the underlying partitioning statistics of dPCR. Using simulations and real-world environmental data, we show that (log-)normality assumptions are violated for dPCR measurements, leading to inaccurate estimates of gene concentrations and underlying biological processes. To enable reliable analyses of environmental dPCR data, we present a dPCR-specific likelihood model that accounts for concentration-dependent measurement noise and non-detects as characteristic of dPCR assays. We demonstrate that this approach overcomes biases in inference from environmental data, such as estimating free-eDNA decay in seawater or pathogen transmission from wastewater monitoring. Our method is implemented in the R packages “dPCRfit” for regression analyses and “EpiSewer” for wastewater surveillance.

## 1 Introduction

The quantification of gene concentrations from environmental samples has become an important technique in environmental science and engineering, with diverse applications in ecology, conservation, and public health. For example, the collection and analysis of environmental DNA (eDNA), i. e. genetic material shed into the environment by living organisms, enables the detailed monitoring of biodiversity and population dynamics [1,2]. By quantifying the abundance of a specific target gene in samples taken at different locations or times, insights into the occurrence, distribution, movements, and interactions of species can be gained [3, 4]. Similarly, in wastewater-based epidemiology, genetic material from pathogens shed by infected individuals into wastewater is sampled at municipal treatment plants to track infection dynamics in human populations without reliance on clinical testing [5,6]. This has enabled the cost-effective monitoring of community-level pathogen spread [7–12] and the estimation of epidemiological parameters, such as the effective reproduction number [13].

The most widespread techniques for quantifying genetic targets in a sample are quantitative PCR (qPCR) and digital PCR (dPCR). While qPCR quantification relies on a comparison of replication cycle thresholds with a standard curve established from reference samples, dPCR works by partitioning a sample into thousands of independent nanoscale chambers or droplets before PCR amplification [14]. Using the ratio of positive partitions to the total number of partitions, the concentration in the sample can be measured [15]. Due to its independence from standard curves, greater robustness and repeatability, improved handling of inhibition, and increased potential for multiplexing, dPCR is becoming an increasingly popular method for quantifying molecular targets [16–18].

To analyze gene concentrations measured in environmental samples from different locations or points in time, various statistical approaches such as non-parametric smoothing [19], correlation analysis [20–22], linear regression [23–25], generalized linear models [26, 27] and latent variable models [28–30] have been used. Although appropriate statistical models exist for the number of positive partitions in a dPCR assay [15], many approaches directly analyze measured concentrations, treating them as normally [19–22, 25, 27, 28] or log-normally [9, 23, 24, 26, 29] distributed. However, such assumptions do not reflect the statistical properties of dPCR assays, which can affect data analysis in several ways. For example, the measurement error in dPCR data increases in a non-linear fashion as sample concentrations become lower [31]. This can lead to unreliable inferences due to heteroskedasticity in ordinary least squares regression or Pearson correlation coefficients [32]. In addition, dPCR measurements may result in non-detects, corresponding to a measured concentration of zero. However, when modeling dPCR data as log-normally distributed, such zero measurements must be removed from the analysis, biasing concentration estimates upwards. These problems are particularly pronounced when analyzing dPCR data with limited sample size and low concentrations, such as estimating species prevalence from a few sampled sites, or monitoring pathogen concentrations at the start of an epidemic wave from weekly wastewater samples.

In this study, we develop an accurate statistical model for analyzing concentration measurements from dPCR. Using established theory of partitioning statistics in dPCR, we derive estimators for the coefficient of variation and the probability of non-detection given the true concentration in a sample. Unlike previous work, we account for noise through extraction, preprocessing, and other sources of variation, which can be substantial in the case of environmental samples. We then propose a likelihood function for estimating gene concentrations and underlying biological process characteristics from reported dPCR measurements. This offers a practical and interpretable alternative to explicitly modeling the number of positive partitions in an assay [15], which is often unavailable from publicly reported data. We demonstrate that, compared to normal and log-normal models, our method achieves more reliable and precise estimates of i) decay rates for marine species free-eDNA in seawater and ii) infection dynamics of respiratory viruses from wastewater.

## 2 Materials and Methods

### 2.1 Partitioning statistics in digital PCR

In digital PCR, the reaction mixture is split uniformly into *m* independent partitions, where *m* is often in the order of tens of thousands. The number of target molecules in each partition is assumed to follow a Poisson process [33], i. e. partition *j* ∈ {1,…, *m*} will contain *N*_*j*_ ∼ Pois(*λc*) molecules. Here, *λ* is the expected concentration in the analyzed sample, and *c* is a “conversion factor” that expresses the expected number of gene copies per partition for one concentration unit of the original sample. This factor can be expressed as a product of the partition volume *v* and a scaling factor *s*, summarizing concentration differences due to preprocessing (e. g. extraction, adding of reagents, dilution, and suboptimal recovery efficiency [34]). During PCR, a fluorescent signal is generated if one or several gene copies are present in a partition, leading to a binary result (positive or negative) for each partition. Since the reaction in each partition is regarded as independent, the total number of positive partitions *Y* is binomially distributed with

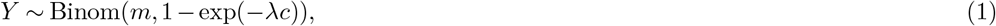

where 1 − exp(−*λc*) is the probability for at least one gene copy being present in a partition.

Under this model, Dorazio and Hunter [15] show that the maximum likelihood estimate for the concentration *λ* given *Y* = *y* observed positive partitions is

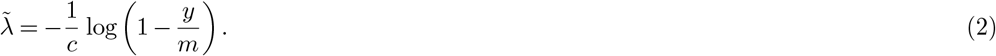

It is furthermore common to run *n* technical replicates *i* ∈ {1,…, *n*} of the same sample. Replicate measurements can then be combined by applying the maximum likelihood estimator to the pooled partition counts or by computing the arithmetic mean of individual estimates (see Supporting Information A for a comparison of both approaches). In the following, we assume an equal number of partitions *m*_*i*_ = *m* for all technical replicates, in which case our results are valid for both approaches. Naturally, by letting *n* = 1, our model also applies to single measurements.

In environmental surveillance, studies often report the maximum likelihood estimate 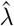 for the target concentration *λ* rather than the number of positive partitions *y*. Here we develop a suitable model for 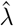 by deriving the coefficient of variation and probability of non-detection of dPCR measurements and proposing an appropriate likelihood function.

### 2.2 Coefficient of variation

We now regard the dPCR measurement 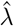 as a random variable 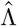. Assuming that *λ* is the true expected sample concentration, and using the Binomial model of positive partition counts *Y*, it can be shown that the asymptotic variance of 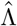 is

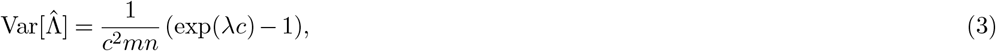

see Supporting Information B.1 for details. Using the fact that 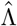 is an asymptotically unbiased estimator of *λ* [33], we can estimate the coefficient of variation (CV) of the measurements as

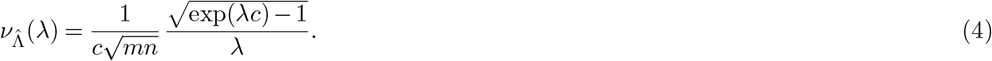

Eq. (4) implies that the CV of measurements is concentration-dependent, which can lead to heteroskedasticity in linear regression even if measurements are log-transformed [32]. However, this estimate assumes that the concentration in the PCR is exactly proportional to the original concentration of interest *λ*. To account for extraction and preprocessing noise and other sources of variation, we extend our model by defining the unscaled concentration in the PCR as a random variable Λ_pre_ that follows a parametric distribution with mean *λ* and coefficient of variation *ν*_pre_ *>* 0. Here, the parameter *ν*_pre_ describes random variation before the PCR. As shown in Supporting Information B.2, the variance of 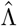 under this extended model is

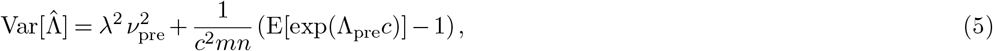

where *E*[exp(Λ_pre_*c*)] can be obtained from the moment-generating function (MGF) of Λ_pre_. For example, if Λ_pre_ is gamma distributed, then

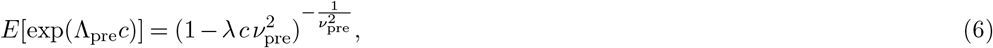

yielding a coefficient of variation

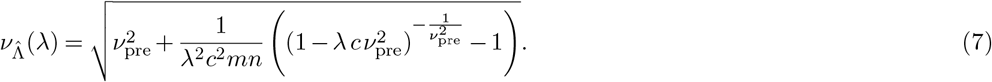

In Supporting Information B.2 we also provide an approximation for the case where Λ_pre_ is log-normally distributed, and use a Taylor series expansion to approximate 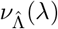 if only its mean and variance are known.

### 2.3 Probability of non-detection

A dPCR run can have zero positive partitions even for non-zero concentrations. If this happens for all technical replicates, the target is not detected, and the maximum likelihood estimate for the concentration is 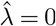. Without accounting for pre-PCR variation, the probability for such a non-detection follows directly from the binomial model in Eq. (1), i. e.

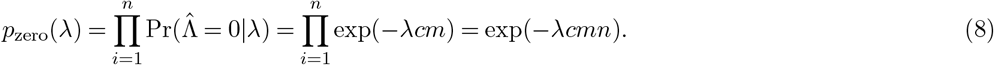

This probability is closely related to the Limit of Detection (LoD) of an assay, which is typically defined as the lowest concentration *λ*_LOD_ at which detection is feasible with sufficient confidence *α* [35]. Leaving the possibility of false positive partitions aside, we note that *α* = 1 − *p*_zero_(*λ*_LOD_) = 1 − exp(−*λ*_LOD_*cmn*), hence *λ*_LOD_ sufficiently determines the non-detection probability under Eq. (8) for arbitrary concentrations (see Supporting Information C.1 for further discussion).

To account for pre-PCR variation as in Section 2.2, we compute the marginal probability of non-detection, which can be expressed as

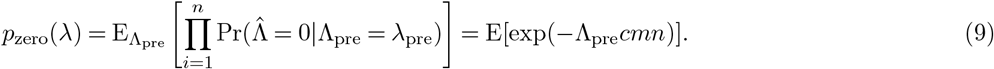

Similar to Eq. (5), the expectation E[exp(−Λ_pre_*cmn*)] can be obtained using the MGF of Λ_pre_, and we provide solutions for the case where Λ_pre_ is gamma or log-normally distributed in Supporting Information C.2.

### 2.4 Likelihood function

Based on the concentration-dependent noise and probability of non-detection in digital PCR as derived above, we propose the following likelihood for a dPCR measurement 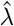:

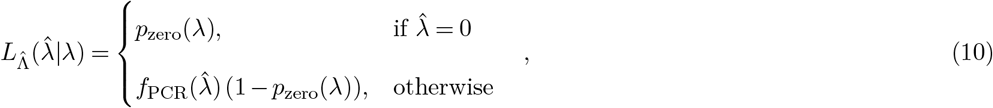

where *p*_zero_(*λ*) is the probability of non-detection as defined in Eq. (9) and *f*_PCR_ is the density function of a positive continuous probability distribution with conditional mean

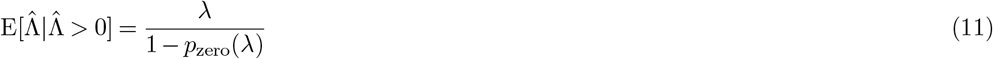

and conditional coefficient of variation

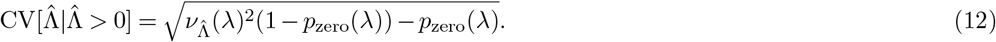

That is, *f*_PCR_ models the distribution of non-zero measurements by adjusting the expected concentration *λ* and the coefficient of variation 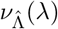 (see Eq. (7)) for the conditioning on a non-zero outcome (see Supporting Information F.1 for details). The likelihood defined in Eq. (10) corresponds to a hurdle model in which concentrations can be measured as zero with a certain probability and otherwise follow the specified continuous probability distribution *f*_PCR_. We note that while 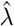 is a real number, its distribution is not continuous due to the underlying count nature of the digital PCR. Therefore, the use of a continuous distribution for *f*_PCR_(*x*) is an approximation of the true distribution, with the advantage of being computationally more efficient than a back-calculated binomial likelihood and having an unbiased mean even if the assumed size or number of partitions is unknown or misspecified (see Supporting Information F.2). We here choose a gamma distribution for *f*_PCR_(*x*), as suggested by a simulation-based comparison of different continuous distributions (see Supporting Information F.3).

### 2.5 Inference from dPCR measurements

Given a set of dPCR measurements 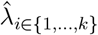 from *k* samples, we can use the likelihood 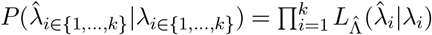 as proposed in Eq. (10) to estimate the underlying concentrations *λ*_*i*∈{1,…,*k*}_ via Bayesian inference. Specifically, given a concentration model *λ*_*i*_ = *f* (*i, θ*) with parameters *θ* and corresponding prior *P* (*θ*), we can sample from the posterior distribution 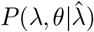 via Markov Chain Monte Carlo (MCMC). In this study, we implemented a generalized linear model (GLM) for *f* (*i, θ*), allowing us to estimate the association of covariates with the target concentration under different link functions. Moreover, as an example of a domain-specific model, we also implemented a model of pathogen concentrations in municipal wastewater for *f* (*i, θ*). Both models are implemented in the probabilistic programming language “stan” [36] and can be fitted through an interface in the R programming language (see Supporting Information G.1 for details on the estimation). We provide a dedicated R package for the GLM (“dPCRfit”) [37] and for the wastewater model (“EpiSewer”) [38].

To apply Eq. (10), we in principle require knowledge of the PCR parameters (number of partitions *m*, scaling factor *s*, partition volume *v*) and the pre-PCR coefficient of variation *ν*_pre_. While these quantities are known in a typical laboratory setting, they may be unknown in other settings, e. g. when modeling publicly reported concentration measurements without sufficient metadata. In this case, it is possible to provide broad priors for these parameters and estimate them jointly with the concentrations *λ*. We provide an example of such a joint estimation using only measurement data when applying our model to wastewater concentration measurements.

### 2.6 Model validation

We used simulations of positive partition counts according to Eq. (1) under different types of pre-PCR noise to validate our approximations for the CV and the probability of non-detection (Supporting Information D). We compared the theoretical predictions for the CV and the probability of non-detection with empirical data from ongoing wastewater surveillance by Eawag, the Swiss Federal Institute of Aquatic Science and Technology (Dübendorf, Switzerland). For this, we used dPCR measurements for Influenza A virus (IAV), Influenza B virus (IBV), Respiratory syncytial virus (RSV), and SARS-CoV-2 of samples from 14 wastewater treatment plants in Switzerland from July 10, 2023 to March 5, 2024 obtained on the Naica Crystal Digital PCR Platform with Sapphire Chips (Stilla Technologies, France) using the laboratory methods described in Nadeau et al. [39]. The empirical CV was estimated based on the bias-corrected empirical standard deviation of two technical replicates divided by their empirical mean. The empirical probability of non-detection was estimated as the percentage of second replicates with a result of zero among all replicate pairs where the result of the first replicate falls within a moving window of 1 gc*/*mL width (Supporting Information E). To validate our proposed likelihood, we estimated concentrations from simulated measurements under the positive partition count model described in Section 2.1 for concentrations of 1, 3, and 5 gc*/*mL, assuming a scaling factor of *s* = 30, a partition volume of *v* = 0.519 nL, and *m* = 25000 partitions per measurement (Supporting Information G.2).

### 2.7 Application to eDNA-based biomonitoring

We conducted a reanalysis of data from an aquarium study by Scriver et al. [27], which tested a direct droplet dPCR assay for eDNA-based marine species monitoring. In the study, the abundance of free-eDNA of the brown bryozoan “Bugula neritina” in aquarium seawater was measured by concentrations of the Cytochrome c oxidase subunit 1 (COI) gene at different points in time up to 72 hours after removal of the organism from the aquarium. The speed of decline in detectable gene concentrations was then estimated using an exponential decay model. We here applied a GLM with logarithmic link using the “dPCRfit” package to model the exponential decay in free-eDNA, measured in terms of the half-life period (see Supporting Information H for details). In the original analysis, the authors used the averages of measurements across three tanks with low, medium, and high initial species biomass. Here, we instead applied our GLM to the individual measurements, and additionally stratified our analysis across tanks. We then compared the consistency of eDNA half-life estimates between tanks and with the original result by Scriver et al. when using a normal, log-normal, or dPCR-specific likelihood.

### 2.8 Application to wastewater-based epidemiology

We combined our dPCR-specific likelihood function with a wastewater model implemented in the R package “EpiSewer” [38] to analyze real-world duplicate dPCR measurements of Influenza A virus from the wastewater treatment plants of Zurich and Geneva, Switzerland, during the 2022/23 Influenza season. We used discretized estimates from the literature to characterize the generation time distribution (shifted Gamma distribution, mean = 2.6, sd = 1.5 [40, 41]) and the shedding load distribution since time of infection (Gamma distribution, mean = 2.49, sd = 1 [42]) of Influenza A virus. Based on a comparison with confirmed case numbers, we assumed an average total shedding load per case of 1 × 10^11^ gc. We provided broad priors for the parameters of the dPCR assay (*c* and *m*) and estimated these jointly from the measurement data (Supporting Information I.1).

We applied the model to longitudinal wastewater measurements to estimate the viral load in wastewater and the effective reproduction number *R*_*t*_ over time. We compared estimates obtained under a normal, log-normal, and dPCR-specific likelihood. We also produced short-term forecasts of concentrations up to one week ahead (assuming continued transmission dynamics) and assessed forecast performance using the continuous ranked probability score (CRPS) [43]; see Supporting Information I.2 for details.

## 3 Results

### 3.1 Coefficient of variation

The coefficient of variation (CV) of measurements from dPCR depends on the target concentration in the sample. As shown in Figure 1A, the CV is approximately constant for high concentrations but increases exponentially as the concentration approaches zero. It also increases when concentrations become extremely high, such that the share of positive partitions approaches one (Figure S5) [31]. The presence of pre-PCR variation further increases the CV of measurements, as shown for gamma distributed pre-PCR noise in Figure 1. Apart from extremely high concentrations, we find that the CV is almost identical for log-normally and gamma-distributed pre-PCR noise and can be well approximated using a Taylor series expansion (Figures S4–S5). The averaging of *n* technical replicates scales the CV by a factor of 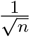. For example, absent pre-PCR noise, the pooling of two wells or, equivalently, the use of an average of two technical replicates reduces the CV by ≈ 30%. Our theoretical CV model also agreed well with empirical CV estimates, based on technical replicates of dPCR measurements in wastewater samples from Switzerland (Figures 1A and S10). For Influenza A virus, we found a median absolute deviation of 4.64 percentage points between the theoretical curve predicted by Eq. (4) and the locally estimated scatterplot smoothing (LOESS) curve for empirical estimates. Since the empirical CV is based on technical replicates of the same preprocessed sample, we could not assess the effect of pre-PCR noise as predicted by Eq. (7).

**Fig 1.**
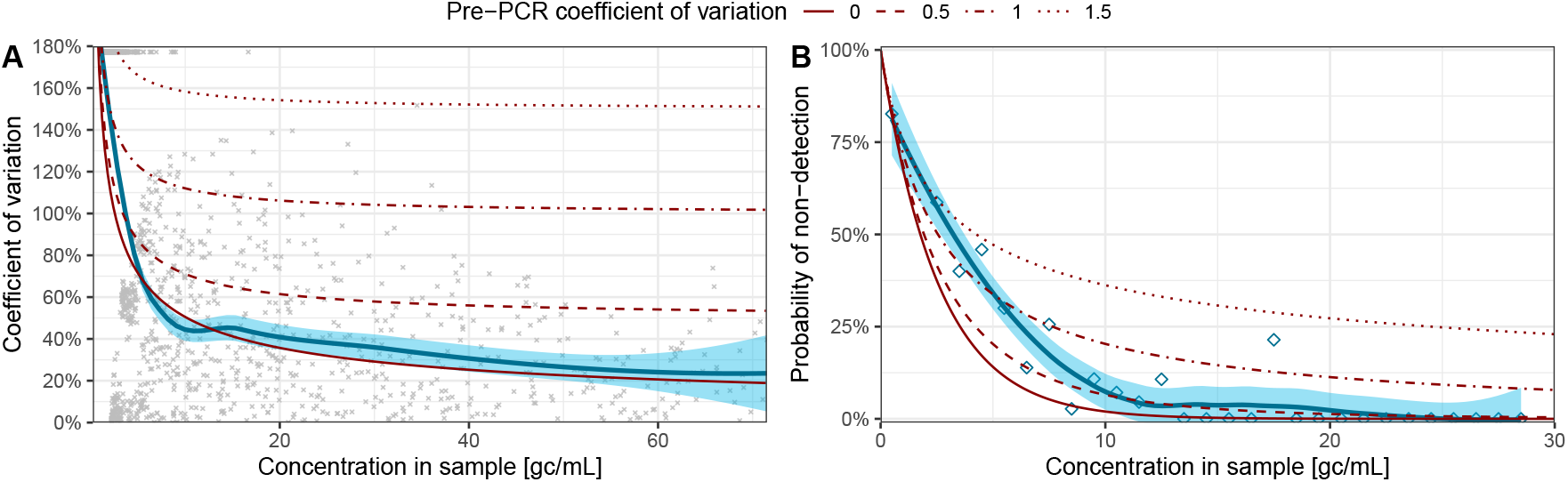
Relationship of the sample concentration with the coefficient of variation and probability of non-detection in digital PCR. Shown is (A) the coefficient of variation (CV) and (B) probability of non-detection (*p*_0_) for a single digital PCR measurement, as a function of the sample concentration *λ*. Empirical estimates of the CV (dots) and *p*_0_ (diamonds) were computed from duplicate dPCR measurements of Influenza A virus from 14 wastewater treatment plants in Switzerland (Naica Crystal Digital PCR Platform, scaling factor *s* = 30, partition volume *v* = 0.519 nL, average partition number *m* = 22665). Blue lines with 95% uncertainty intervals show LOESS-based kernel regressions of the empirical estimates, indicating the relationship with the sample concentration *λ*, respectively. Red lines show the CV and *p*_0_ as predicted by Eq. (7) and Eq. (9), for different strengths of the pre-PCR coefficient of variation *ν*_pre_ under gamma distributed pre-PCR noise (dashed and dotted lines). As the empirical estimates are based on technical replicates of the same sample, they do not include pre-PCR variation and therefore correspond to *ν*_pre_ = 0.

### 3.2 Probability of non-detection

The probability of non-detection describes the chance of zero measurements despite the presence of the target in the sample. Such “false negatives” are problematic when modeling dPCR measurements via strictly positive likelihoods such as a log-normal distribution, as they assign zero probability to such measurements. If concentrations are sufficiently low, non-detects will however occur. For the PCR parameters described in Section 2.6, Eq. (8) predicts that even without pre-PCR noise, there is an 11.5% probability that a concentration of *λ* = 5 gc*/*mL will be measured as zero. For gamma-distributed pre-PCR noise with a CV of *ν*_pre_ = 0.5, the non-detection probability increases by another 5.5 percentage points (Figure 1B). Importantly, we find that the non-detection probability curve changes significantly when we model the pre-PCR noise as log-normal instead of gamma-distributed (Figure S9), with gamma-distributed noise leading to higher non-detection probabilities. Moreover, the probability of non-detection decreases with the number of partitions and the number of replicates (Eq. (8)). For example, a doubling of the number of replicates has the same effect on the probability of non-detection as a doubling of the concentration. We compared our model for the probability of non-detection with empirical estimates based on the same real-world measurements as in Section 3.1 (Figures 1B and S10). Although the empirical estimates rely on a moving window approach and are therefore less accurate, we find a general agreement between the theoretical curves predicted by Eq. (8) and the LOESS curve of the empirical estimates (median absolute deviation: 3.40 percentage points for Influenza A).

### 3.3 Application to eDNA-based biomonitoring

We used a generalized linear model with our dPCR-specific likelihood function to estimate the half-life of free-eDNA of a marine species in an aquarium seawater experiment by Scriver et al. [27]. In the study, concentrations of the Cytochrome c oxidase subunit 1 (COI) gene of the brown bryozoan “Bugula neritina” were measured in different aquariums with low, medium, and high biomass of the species, using samples taken 0, 4, 8, 24, 48, and 72 hours after removal of the organism from the aquarium. Scriver et al. estimated the exponential decay rate of detectable free-eDNA from the averages of all measurements at each point in time. We reproduced this analysis by fitting a GLM with normal likelihood to the averaged measurements. However, when stratifying the analysis to individual aquariums, i. e. each with smaller sample size, this averaging approach yielded inconsistent results (Figure 2A). When fitting to individual samples using a normal or log-normal likelihood, results remained inconsistent. Specifically, the normal likelihood yielded half-life estimates much lower than the original estimate by Scriver et al. Under the log-normal likelihood, half-life estimates were higher than the original estimate and highly uncertain. In contrast, our dPCR-specific likelihood yielded precise and consistent results across aquariums. The median estimates of the free-eDNA half-life were 19.51 (95% CrI 12.07–45.24), 12.63 (95% CrI 8.95–20.13), and 15.50 (95% CrI 11.17–25.87) hours for low, medium, and high initial biomass, and 18.75 (95% CrI 14.36–26.85) hours for all aquariums combined, which agrees well with the original estimate by Scriver et al. When comparing median concentrations as predicted by the models, we found that predictions under the dPCR-specific likelihood were generally between those under the normal and log-normal likelihood (Figure 2B).

**Fig 2.**
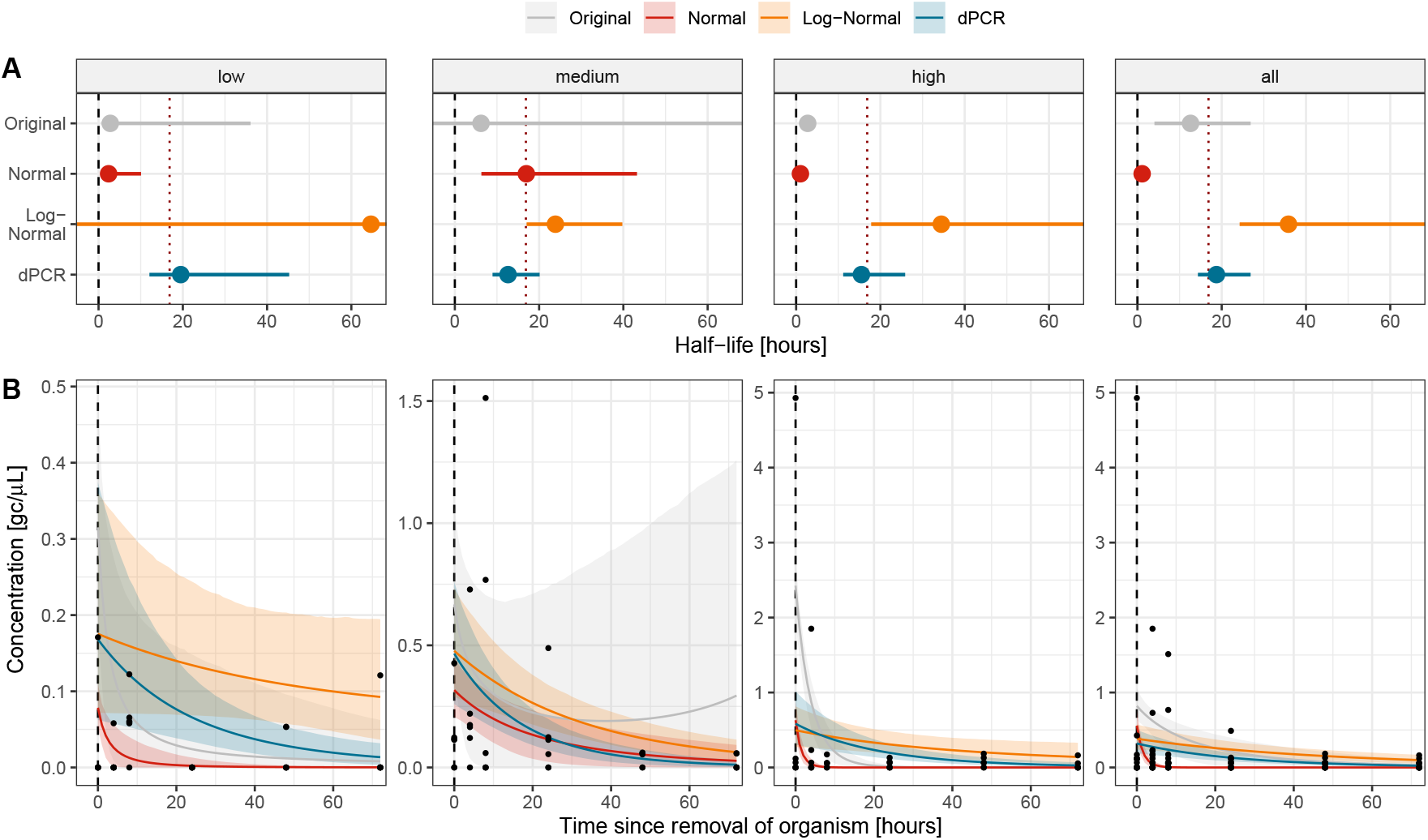
Modeling the exponential decay of eDNA of a marine species in aquarium seawater. Estimated half-life period for free-eDNA of Bugula neritina in aquarium seawater based on longitudinal dPCR measurements after removal of the organism by Scriver et al. [27]. Shown are median estimates (dots) with 95% credible intervals (bars) from a generalized linear model (GLM) applied to data from an aquarium with low, medium, and high initial biomass, respectively, and from all aquariums combined. The GLM was applied to averages of measurements as in the original analysis by Scriver et al. (grey) and to individual measurements using a normal (red), log-normal (orange), and dPCR-specific (blue) likelihood function. Red vertical lines show the original estimate reported by Scriver et al. (B) Measured concentrations (dots) and model predictions using the different likelihoods (colored lines) with 95% credible intervals (shaded areas).

### 3.4 Application to wastewater-based epidemiology

The use of a dPCR-specific likelihood can improve wastewater-based surveillance of pathogen transmission. Using the normal, log-normal, and dPCR-specific likelihood, we computed estimates and forecasts of concentrations, viral loads, and effective reproduction number *R*_*t*_ values from dPCR measurements of Influenza A virus at the municipal treatment plant of Zurich, Switzerland (Figure 3). Under the normal likelihood, measurements are modeled to have constant variance, leading to an overestimation of measurement noise at low concentrations. For example, before Dec 1, 2022, the 80% credible intervals (CrIs) of measurements predicted by the normal likelihood covered already 98.5% of observations, suggesting a miscalibration of the model (Figure 3A). The concentration curve predicted by the normal likelihood model was also highly volatile, with the median estimated concentration falling within the previous 95% CrIs on only 56.0% of days. This also resulted in volatile estimates of the viral load (Figure 3B) and the effective reproduction number (Figure 3C), likely due to overfitting of the model at the peak of the seasonal wave. *R*_*t*_ estimates at the start of the seasonal wave were highly uncertain (Figure 3C). Under the log-normal likelihood, measurements are assumed to have a constant coefficient of variation, which is approximately valid at high concentrations. This resulted in more similar estimates by the log-normal and dPCR-specific likelihood models. Nevertheless, measurement noise tended to be underestimated for low concentrations (80% CrIs covered only 59.7% of observations before Dec 1, 2022) and overestimated for high concentrations (80% CrIs covered 100% of observations after Dec 1, 2022) by the log-normal model. As a result, the estimated and forecasted load curve and *R*_*t*_ estimates were more uncertain than under the dPCR-specific likelihood. By modeling the concentration-dependent coefficient of variation, the dPCR-specific likelihood model achieved similar coverage at low and high concentrations (80% CrIs covered 88.0% of observations before Dec 1, 2022, and 84.2% after Dec 1, 2022) and estimated viral load and *R*_*t*_ values with reduced uncertainty. It also achieved more accurate forecasts of measurements after Dec 26, 2022 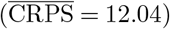 than the normal 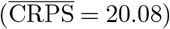 and log-normal likelihood 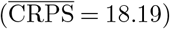. In Figures S16–S20, we show additional results from fitting the models on different date ranges of measurements at the treatment plants of Zurich and Geneva, Switzerland.

**Fig 3.**
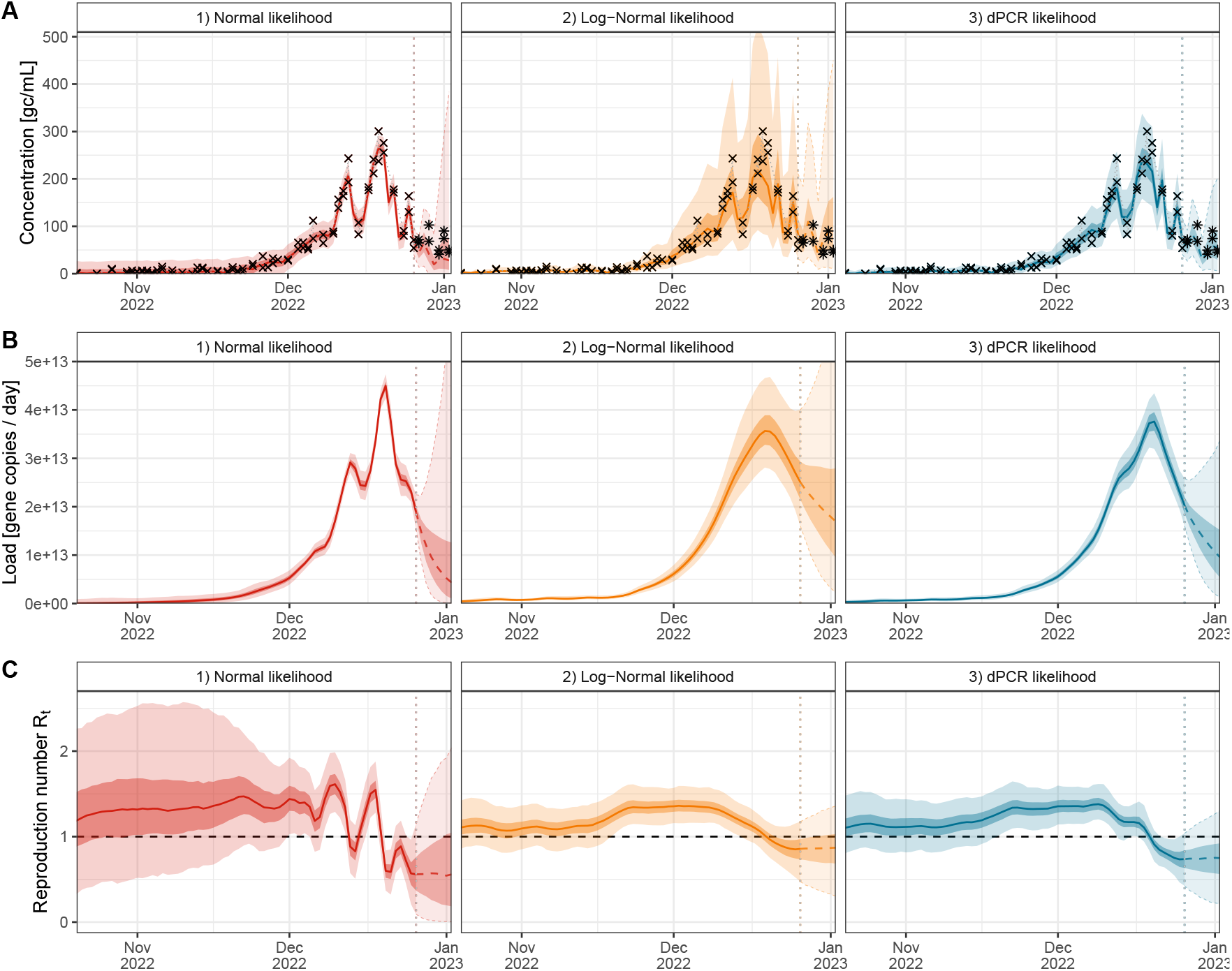
Comparison of inference and short-term forecasts from wastewater dPCR measurements using different likelihood functions: Shown are the results of fitting an epidemiological wastewater model (EpiSewer) to duplicate dPCR measurements of Influenza A virus concentrations at the municipal treatment plant of Zurich, Switzerland (Oct 20 – Dec 26, 2022) using either 1) a normal likelihood (red), 2) a log-normal likelihood (orange), and 3) a dPCR-specific likelihood (blue) for the observations. Each panel shows the median (lines) and 50% and 95% credible intervals (strong and weakly shaded areas) of A) posterior predictive distributions for dPCR measurements, B) estimated viral loads in wastewater over time, and C) the estimated effective reproduction number over time. Estimates shown beyond Dec 26, 2022 (vertical dotted line) are forecasted based on a continuation of the latest transmission dynamics. Panel A also shows the true observed (crosses) and future (stars) dPCR measurements.

When fitting the dPCR-specific model to wastewater measurements, we assumed that the precise characteristics of the wastewater assay are unknown and only assigned broad priors for the number of total partitions *m* and the conversion factor *c*. Despite the additional parameter uncertainty, this yielded highly similar estimates as a fully informed model where *m* and *c* are fixed to their true values (Figure S21). Moreover, additional insights about the measurement process can be gained from the estimated laboratory parameters. For example, the posterior distribution for *ν*_pre_ had a median of 17.0% (95% credible interval: 5.3% – 28.1%) and differed clearly from its prior (Figure S15). As this parameter describes the variation in concentrations before the PCR, it allows us to distinguish the technical noise of dPCR from other sources of variation. For instance, using the median estimate for *ν*_pre_ and for the given values of *m* and *c*, the overall CV at a concentration of 100 gc*/*mL is 22.9%, which is 7.8 percentage points higher than the theoretical minimum CV that would be achieved without preprocessing noise or other unexplained variation (15.1%).

## 4 Discussion

In this work, we have developed a model for the statistical analysis of dPCR concentration measurements. Our model reflects two important properties of dPCR data that violate standard assumptions in regression analysis. First, measurement noise in dPCR is concentration-dependent, with the coefficient of variation increasing exponentially as concentrations become small. When not accounted for, this heteroskedasticity leads to unreliable estimates especially at small sample sizes. Second, even with the target gene present in a sample, there can be a significant probability of non-detection, resulting in zero measurements. These data must be removed or imputed when modeling measurements as log-normally distributed, leading to biased estimates. Our likelihood function avoids these problems by modeling both concentration-dependent measurement noise and the probability of zero measurements while accounting for pre-PCR variation, e. g. from sample extraction and preprocessing.

Using real-world examples of dPCR measurements, we have shown that our model achieves more precise and reliable estimates from dPCR data than simpler likelihoods currently used in practice, such as normal or log-normal distributions. This can impact inferences about underlying biological processes, such as eDNA decay in biomonitoring or pathogen transmission dynamics in wastewater-based epidemiology. We argue that a log-normal distribution with a constant coefficient of variation (or, equivalently, modeling log-transformed measurements as normally distributed) may provide a sufficient approximation, but only if all concentrations are sufficiently high, e. g. above the Limit of Quantification (LoQ), which is operationally defined as a concentration below which the CV surpasses a fixed threshold such as 30% [35]. When concentrations are lower, it is important to model a concentration-dependent coefficient of variation and the probability of zero measurements to avoid biased or highly uncertain estimates.

Our findings also have broader implications for data analysis in dPCR-based environmental surveillance. In particular, we argue that statistical models should be applied directly to measured concentrations rather than to derived indicators. For example, in wastewater surveillance, measured pathogen concentrations are often normalized by flow rates (“loads”) or fecal biomarker concentrations, such as crAssphage or PMMoV [44, 45]. However, these normalized quantities are not ideal as direct inputs to wastewater models since their measurement distribution depends on the normalization factor. For example, depending on the flow rate, the same “load” may correspond to vastly different concentrations, with different measurement noise and probabilities of non-detection. The same applies to concentrations normalized by fecal markers, with the added complication that marker concentrations are themselves subject to measurement noise. Therefore, we recommend using non-normalized concentrations as observations and incorporating normalization factors into the model through a latent variable approach, as e. g. done in the R package “EpiSewer” [38].

Moreover, for modeling dPCR measurements at low concentrations, the likelihood proposed in this work should be more appropriate than censoring approaches in which zero measurements or measurements below the LoD are replaced with a predefined value [24, 46]. This is because censoring uses a fixed threshold, while the probability of non-detection gradually increases as the concentration approaches zero. Furthermore, censoring applies to the measurements, which are subject to noise, not to the true concentration in the sample. Similar arguments apply to the LoQ, considering the gradual increase of the CV as concentrations approach zero and the fact that the CV depends on the true sample concentration, not the measurement. We thus argue that it is better to model the increase in CV at low concentrations than to classify measurements by a predefined threshold. Indeed, removing measurements below the LoQ from statistical analyses may lead to an overestimation of concentrations.

Using Bayesian inference, our dPCR-specific likelihood can be fitted directly to concentration measurements, as are primarily reported by studies and laboratories. This makes our approach widely applicable, even in settings where the number of positive and total partition counts is not reported. We note that in our examples, concentrations were measured in gene copies per volume, but in the case of solids-based extraction, measurements can also have other units, such as gene copies per gram of dry weight [44]. Information about sample dilution and parameters of the PCR, such as partition number and volume, can be integrated by fixing the respective parameters to their values. However, in our application to wastewater-based epidemiology, our model achieved precise estimates even when the PCR parameters were unknown and broad priors were used.

We note a few limitations of our model. First, we have tested our estimators for the CV and probability of non-detection only for log-normally and gamma-distributed pre-PCR noise. In principle, any positive continuous distribution with a well-defined moment generating function (MGF) could be used, as long as a closed-form expression or accurate approximation of the MGF is available. We acknowledge that choosing a pre-PCR noise distribution can be difficult, as this parameter summarizes all unexplained variation in a model. For the CV, we found that the exact distribution of pre-PCR noise is irrelevant for most concentration values and can be well approximated using a Taylor series expansion. For the probability of non-detection, however, we found notable differences between log-normally and gamma-distributed pre-PCR noise and no suitable Taylor series approximation. Second, we assumed that in expectation, the dPCR measurements are equal to the true sample concentration, i. e. we do not account for unknown systematic distortions of dPCR measurements such as from miscalibration. When estimating PCR parameters jointly from the data, we also assumed that these are stable over time, i. e. systematic changes in the laboratory protocol may not be detected without additional information. Third, we did not account for variation in the volume of partitions, which can be an additional but minor source of noise in droplet-based systems [31, 47]. Fourth, our hurdle model assumes that, given a true concentration and the parameters of the PCR assay, the probability of non-detection is stochastically independent of the CV of the measurements. In practice, these quantities are correlated due to the same underlying pre-PCR noise, however, this correlation has limited impact as it only applies to measurements that are non-zero but small. Fifth, when computing the CV and the probability of non-detection for measurements that have been averaged over several replicates, we assume for simplicity that the number of partitions does not vary between the replicates and that all replicates are based on the same preprocessed sample. If the exact total and positive partition numbers are known for each replicate, this information can be used by fitting our model to individual replicate measurements. Finally, since this work focused on the quantification of targets already present in the environment, we assumed that the true concentration can be arbitrarily small but is larger than zero. If, instead, the objective is to determine whether a particular target is present or absent, it may also be important to model the probability of false positives [35].

We have implemented our dPCR-specific likelihood in the R package “dPCRfit” [37], providing a Bayesian generalized linear model for regression analyses of dPCR data, e. g. to improve eDNA abundance estimates in biomonitoring [25, 26]. We have also integrated the likelihood as a module in the “EpiSewer” package, e. g. to improve wastewater-based estimates of pathogen transmission rates [38]. Our theoretical results can also benefit assay development and optimization by informing the effect of laboratory parameters on the precision of measurements. For example, we show that pre-PCR noise strongly influences the probability of non-detection, which highlights the importance of including extraction and preprocessing steps when determining the Limit of Detection [34, 35]. While this work has focused on digital PCR, we expect that similar statistical relationships and models can be established for quantitative PCR methods [48]. Overall, we envision that accurate modeling of laboratory and measurement processes will improve environmental surveillance and provide new insights into environmental variability by disentangling it from measurement noise.

## Declarations

### Ethics approval

Ethics approval was not required for this study.

### Author Contributions

**Conceptualization:** AL

**Data curation:** AL

**Formal Analysis:** AL

**Funding acquisition:** TJ, TS

**Investigation:** AL **Methodology:** AL

**Project administration:** AL

**Resources:** TS **Software:** AL

**Supervision:** TJ, TS

**Validation:** AL

**Visualization:** AL

**Writing – original draft:** AL

**Writing – review & editing:** AL, TJ, TS

## Acknowledgments

This work contains a secondary analysis of open wastewater data provided by EAWAG, Swiss Federal Institute of Aquatic Science and Technology. We kindly thank the members of the Wastewater Monitoring Laboratory, including Lea Caduff, Sheena Conforti, Seju Kang, Jolinda de Korne-Elenbaas, Charles Gan, Rachel McLeod, and Melissa Pitton, for their contributions to the development of the underlying digital PCR assays and their application in ongoing wastewater surveillance to produce these data. We thank the Swiss Federal Office of Public Health FOPH for supporting the wastewater surveillance by EAWAG.

## Data availability

All data, code for simulation, and analysis scripts are publicly available at https://github.com/adrian-lison/dPCR-observation-model-study.

## Supporting Information

### A Averaging of dPCR measurements

There exist different approaches to combine replicate measurements of a sample into a single concentration estimate. Dorazio and Hunter [1] describe a maximum likelihood estimator for *λ*

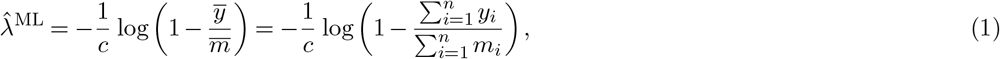

based on replicate measurements *i* ∈ {1,, *n*} of a sample with concentration *λ*, where the replicate runs had *m*_1_, …, *m*_*n*_ total partitions and *y*_1_, …, *y*_*n*_ positive partitions, respectively. This approach is typically used when pooling droplet counts from several wells.

A different approach also used in practice is to compute the arithmetic mean of individual ML estimates for each replicate

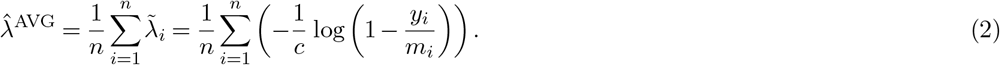

In essence, the ML estimate in Eq. (1) differs from the arithmetic mean of estimates in Eq. (2) in two regards. First, the ML estimate computes an average of the estimated probabilities *p*_1_,, *p*_*n*_ for positive partitions, not an average of the concentrations. Second, this average is weighted by the total number of partitions of each replicate, while the arithmetic mean gives equal weight to all replicates.

To assess the magnitude of these differences, we simulated duplicate dPCR measurements for various concentrations. While the first replicate was assumed to have 25000 total partitions, we simulated the second replicate to have between 5000 and 25000 partitions in order to study the effect of differences in the total number of partitions. We assumed a scaling factor of *s* = 30 and a partition volume of *v* = 0.519 nL. As a first step, we compared the arithmetic mean estimate 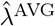 with an unweighted ML estimate

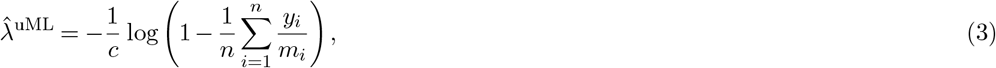

in which each replicate receives the same weight. Thereby, we could isolate the effect of averaging concentrations instead of averaging probabilities. Figure S1 shows how the arithmetic mean of measurements deviates from the unweighted ML estimate. We find that the arithmetic mean has a small positive bias compared to the unweighted ML estimate, as the concentration estimate is a non-linear function of the probability of positive droplets, such that the average of concentrations is not equal to the concentration estimate based on the average probability. The potential deviation increases linearly with the concentration in the PCR, however even for high concentrations of 1000 gc*/*mL, the deviation is considerably less than 1 gc*/*mL.

Next, we compared the unweighted ML estimate defined in Eq. (3) to the weighted ML estimate defined in Eq. (1). We found larger deviations than between the arithmetic mean and unweighted ML estimate, but no systematic bias (Figure S2). Deviations were particularly large in cases where one of the replicates only had few total partitions, such that the corresponding concentration estimate was noisy. In the arithmetic mean and the unweighted ML estimate, such replicates are weighted equally to other replicates. The ML estimate defined in Eq. (1) gives more weight to replicates with more partitions, making it less sensitive to noisy measurements based on few partitions. In practice, however, differences in the total number of partitions larger than 5000 do not occur frequently in dPCR assays. For smaller differences in the total number of partitions, the differences between unweighted and weighted estimates are considerably smaller. Overall, we conclude that the ML estimate of multiple replicates has higher accuracy and precision than the arithmetic mean of individual estimates, but in practice the differences will be small compared to the overall uncertainty of measurements, except in cases where the total number of partitions varies greatly.

**Fig S1.**
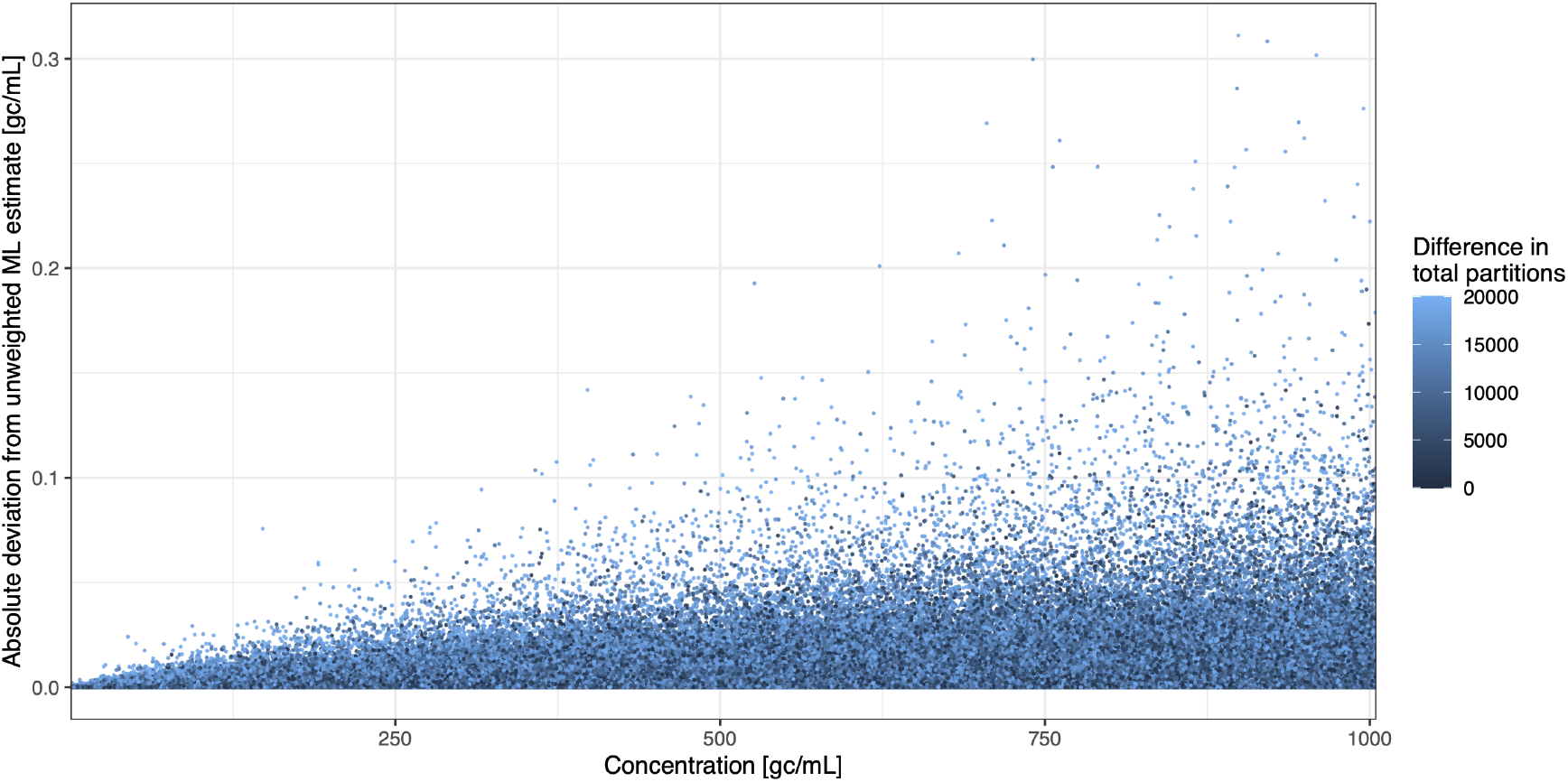
Comparison of arithmetic mean estimate with unweighted ML estimate of concentrations. Shown are deviations of the arithmetic mean of replicate measurements from an unweighted ML estimate for simulated measurements as a function of concentration, and for different degrees of partition number variation between replicates. Simulated measurements assume a scaling factor of *s* = 30 and partition volume of *v* = 0.519 nL.

**Fig S2.**
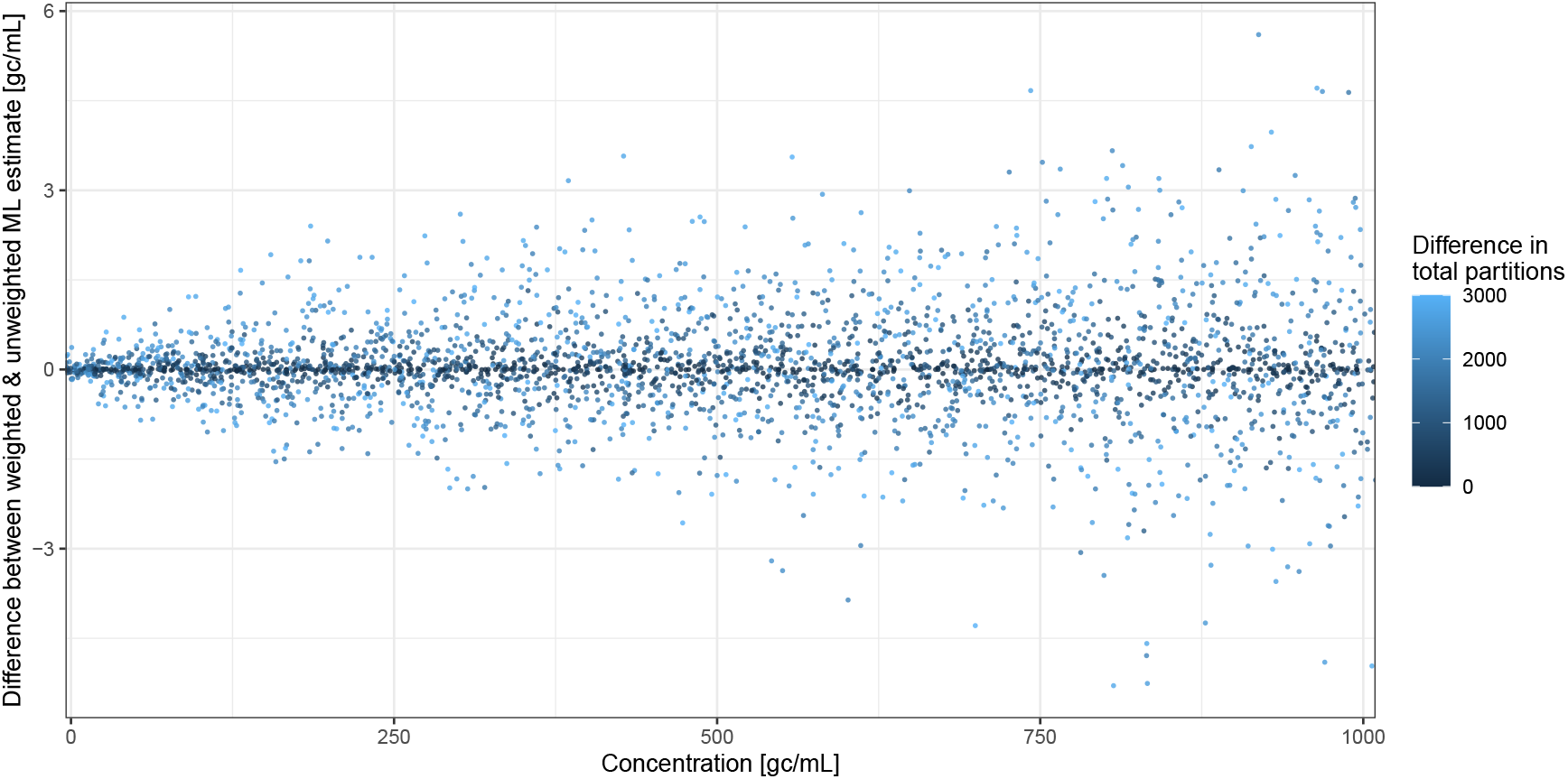
Comparison of unweighted and weighted ML estimates of concentrations. Shown are deviations of the unweighted ML estimate from the weighted ML estimate for simulated measurements as a function of concentration, and for different degrees of partition number variation between replicates. Simulated measurements assume a scaling factor of *s* = 30 and partition volume of *v* = 0.519 nL.

### B Coefficient of variation of concentration estimates

#### B.1 CV without pre-PCR variation

To derive the variance of a single maximum likelihood concentration estimate 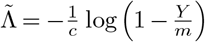 [1], we use the approximation 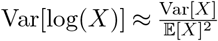 to obtain

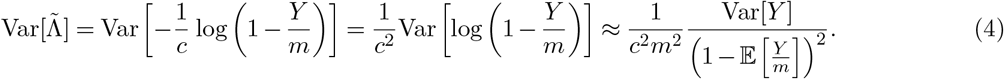

Since *Y* is Binomially distributed with Var[*Y*] = *mp* (1 − *p*) and 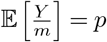, we further get

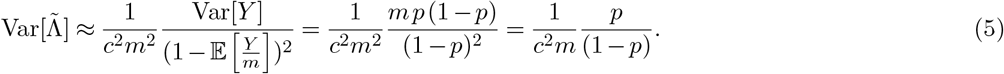

Finally, using the probability for a positive partition *p* = 1 − exp(−*λc*) [1], we obtain

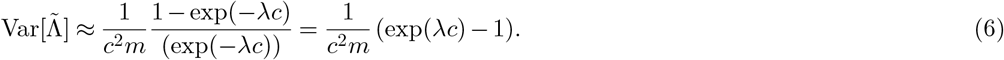

That is, the variance of 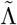 increases exponentially with the concentration in the PCR. This relationship between the variance of 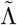 and *λ* can also be obtained through the Fisher information of 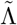 [1].

If we have multiple technical replicates *i* ∈ {1,…, *n*}, the combined maximum likelihood estimate by Dorazio and Hunter as described in Eq. (1) corresponds to a pooled analysis where the *m*_*i*_ partitions from each well are combined into one large assay. Assuming an equal number of partitions *m*_*i*_ = *m* for all replicates, the variance of this estimate is simply

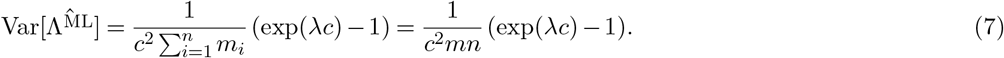

Moreover, if all *m*_*i*_ = *m*, the concentration estimates from individual replicates have identical variances 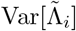, such that the their arithmetic mean as described in Eq. (2) has variance

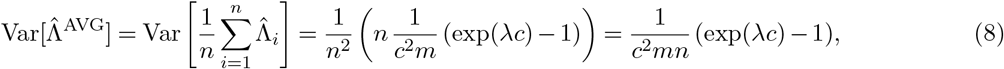

which is identical to the variance of the maximum likelihood estimate in Eq. (7).

Since 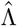 is an asymptotically unbiased estimator of the true concentration [2], i. e.

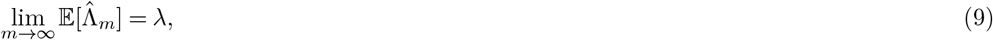

the coefficient of variation of 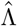 can be estimated as

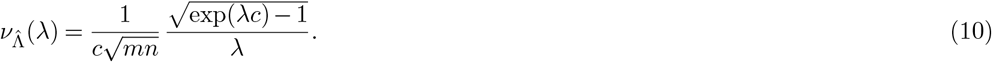

### B.2 CV with pre-PCR variation

As described in the main text, we account for pre-PCR noise by extending the above model such that the concentration in the PCR is *s*Λ_pre_, where Λ_pre_ is a random variable drawn from a parametric distribution with mean *λ* and coefficient of variation 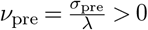. Using the Law of Total Variance, we can express the variance of 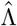 under pre-PCR noise as

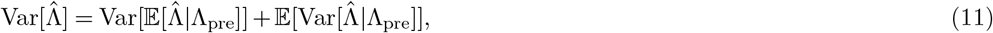

with the “explained” component of the variance

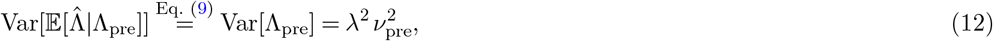

and the “unexplained” component of the variance

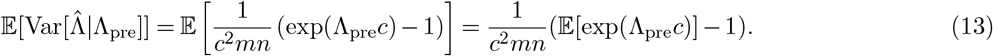

#### B.2.1 Gamma distribution

If we model Λ_pre_ as gamma distributed with parameters 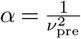 and 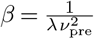, an exact solution for the “unexplained” component of the variance can be obtained. That is, by the MGF of the gamma distribution, we know that

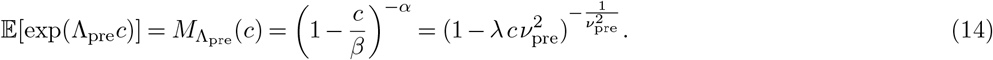

Substituting back, we get the variance

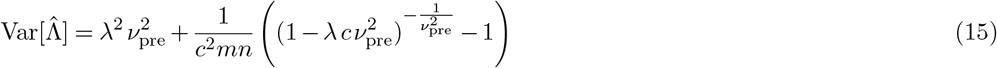

and thus

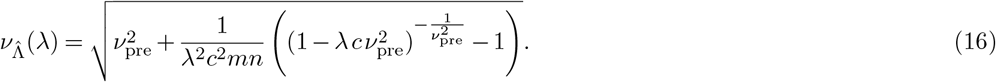

#### B.2.2 Log-normal distribution

We can also model Λ_pre_ as log-normally distributed with parameters 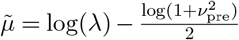 and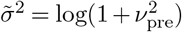. While no closed-form solution for the MGF of the log-normal distribution exists, we can use an approximation [3] to obtain

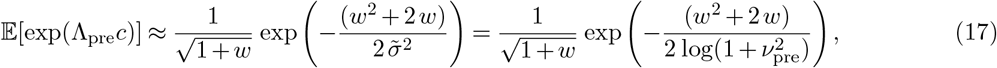

where *w* is an evaluation of the Lambert *W* function [4] with

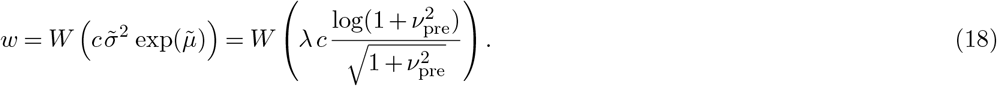

#### B.2.3 Taylor series expansion

If only the mean and coefficient of variation of Λ_pre_ are known, we can approximate *E*[exp(Λ_pre_*c*)] using a Taylor polynomial, i. e.

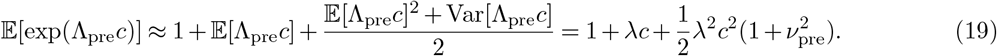

Substituting this back into Eq. (11), we obtain

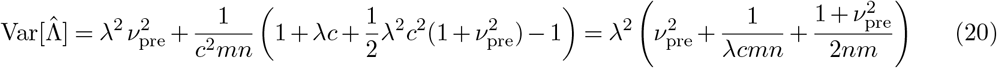

and thus

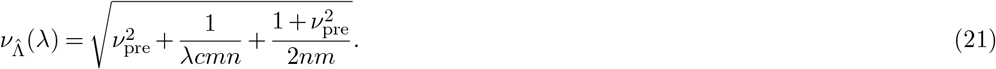

### C Probability of non-detection

#### C.1 Relationship between probability of non-detection and limit of detection

The Limit of Detection (LoD) of an assay is typically defined as the lowest concentration *λ*_LOD_ at which detection is feasible with sufficient confidence *α* [5]. If we leave aside the probability of false positive partitions, this corresponds to the converse of the probability of a zero measurement, i. e. *α* = 1 − *p*_zero_(*λ*_LOD_) = 1 − exp(−*λ*_LOD_*cmn*).

If the specific parameters of an assay are unknown, but the LoD is reported, it may thus be possible to obtain a rough estimate of *cmn* from the reported LoD via

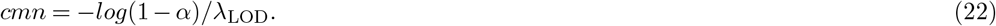

Once an estimate of *cmn* is available, the probability of non-detection can be computed for arbitrary concentrations in the assay. This may be more accurate than censoring measurements below the LoD, as discussed in the main text. On the other hand, as the details of the experiment used to establish the LoD may also be unknown and could violate the above simple relationship, we believe it is generally better to jointly estimate the unknown PCR parameters from the provided non-zero measurements, as described in the Methods.

#### C.2 Probability of non-detection with pre-PCR variation

As described in the main text, the probability of non-detection under pre-PCR variation is

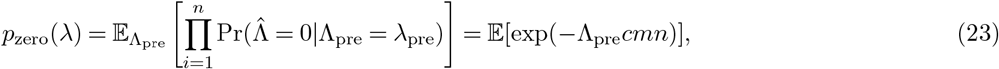

where, as before, Λ_pre_ is drawn from a parametric distribution with mean *λ* and coefficient of variation 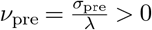. Moreover, 𝔼 [exp(−Λ_pre_*cmn*)] is again an evaluation of the MGF of Λ_pre_. As the Taylor series expansion of Eq. (23) is alternating and its partial sum up to the 2^nd^ degree has a large error, we cannot provide a suitable approximation without further assumptions about Λ_pre_. However, if we assume that Λ_pre_ is gamma distributed, we can again obtain an exact solution using its MGF, i. e.

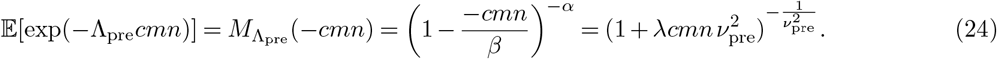

If we instead assume that Λ_pre_ is log-normally distributed, we can use the approximation of its MGF as in Eq. (17), but now with

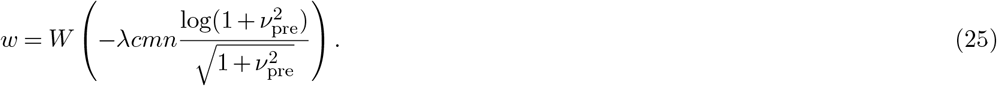

### D Simulation analysis

In the following section, we provide results of simulation analysis in which we compared our analytical approximations for the coefficient of variation and the probability of non-detection with simulated dPCR measurements. For our simulations, we assumed exemplary PCR parameters based on the Naica Crystal Digital PCR Platform with Sapphire Chips (Stilla Technologies, France), a scaling factor of *s* = 30, a partition volume of *v* = 0.519 nL, and *m* = 25000 partitions per measurement, but we also provide a sensitivity analysis of these parameters in Figures S6 and S7. Positive partition counts were simulated under the binomial model described in the main text and transformed into a concentration estimate using the maximum likelihood estimator by Dorazio and Hunter [1], see main text.

#### D.1 Coefficient of variation

Figure S3 shows the coefficient of variation (CV) as a function of the sample concentration under gamma distributed pre-PCR noise. The theoretical prediction for the CV based on Eq. (16) matches the simulated dPCR data well. As can be seen, pre-PCR variation acts as an intercept for the CV function only at higher concentrations, and the CV functions converge as the concentration approaches zero.

Figure S4 shows the coefficient of variation (CV) as a function of the sample concentration under log-normally instead of gamma distributed pre-PCR noise. In addition to the CV of simulated measurements, two theoretical approximations are shown. While dashed lines show an approximation of the log-normal MGF by Asmussen et al. [3] (see Supplement B.2.2), straight lines show the CV as approximated using a Taylor series expansion (see Supplement B.2.3). As can be seen, the approximations yield fairly similar results, however, we find that for high levels of pre-PCR variation, the MGF-based approximation gives slightly lower CV values than the Taylor series-based approximation.

**Fig S3.**
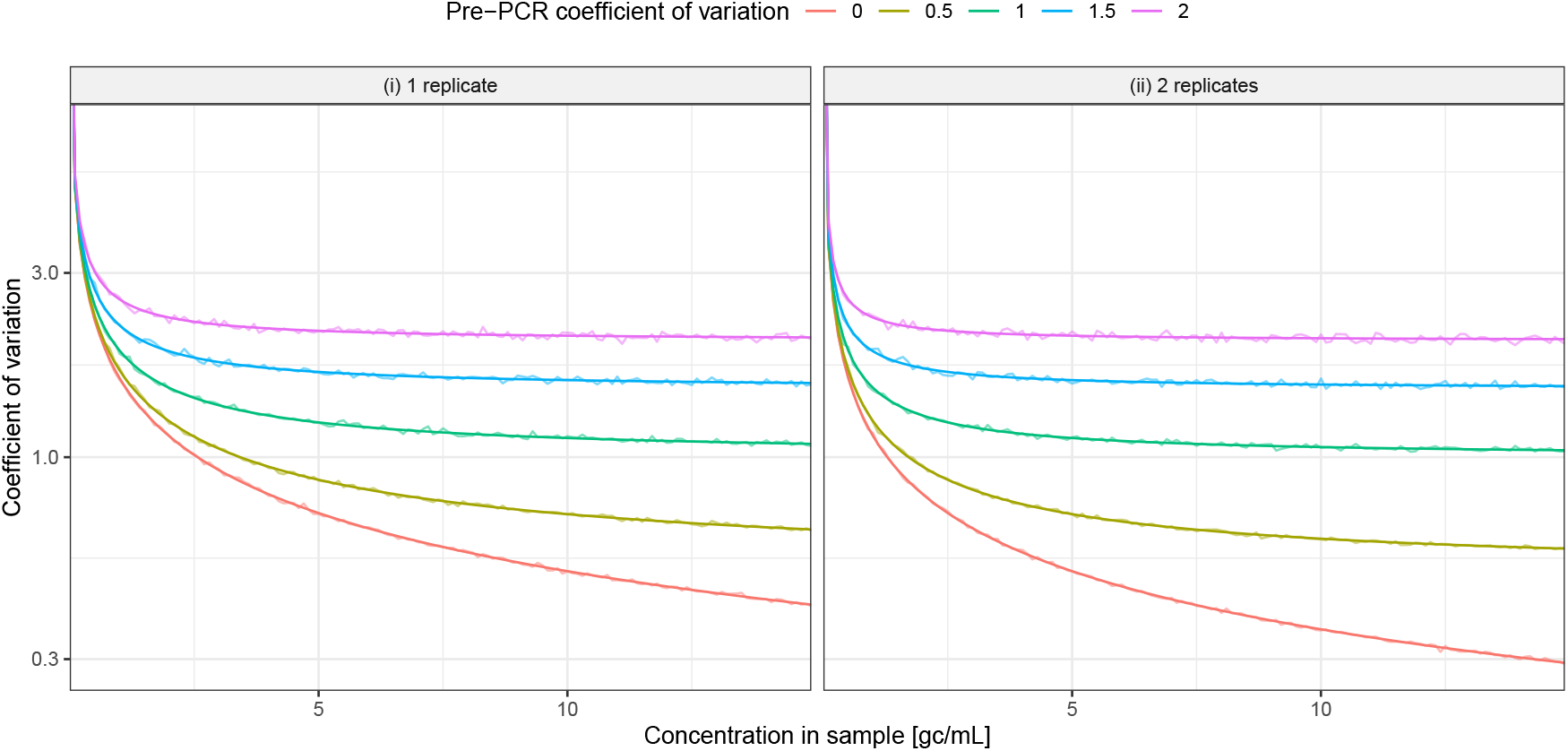
Relationship between concentration and coefficient of variation of dPCR measurements under gamma distributed pre-PCR noise. Shown is the coefficient of variation (CV) of measurements from digital PCR for (i) a single replicate and (ii) the arithmetic mean of two replicates, as a function of the sample concentration *λ* under gamma distributed pre-PCR noise. Jagged lines show the CV of simulated measurements for different concentrations in steps of 0.1 gc*/*mL. Straight lines show the corresponding CV as predicted by Eq. (16). Colors correspond to different strengths of pre-PCR noise, as measured by the pre-PCR coefficient of variation *ν*_pre_.

Figure S5 compares the exact solution for gamma distributed pre-PCR noise with the log-normal MGF approximation and the Taylor series-based approximation across a wide range of concentrations. We note that the upper limit of concentrations shown (900000 gc*/*mL) corresponds to an extreme case where the expected number of gene copies per partition is 15 or higher (in practice, dPCR assays aim for concentrations that correspond to an expected number of gene copies well below 1, which can be achieved via dilution). As the share of positive partitions in the PCR approaches 1, the CV increases considerably. This increase in CV at extreme concentrations is not captured by the Taylor series expansion approximation but by the exact solution under gamma noise and the MGF approximation under log-normal noise. There are small differences in the CV between the gamma case and the log-normal approximation.

**Fig S4.**
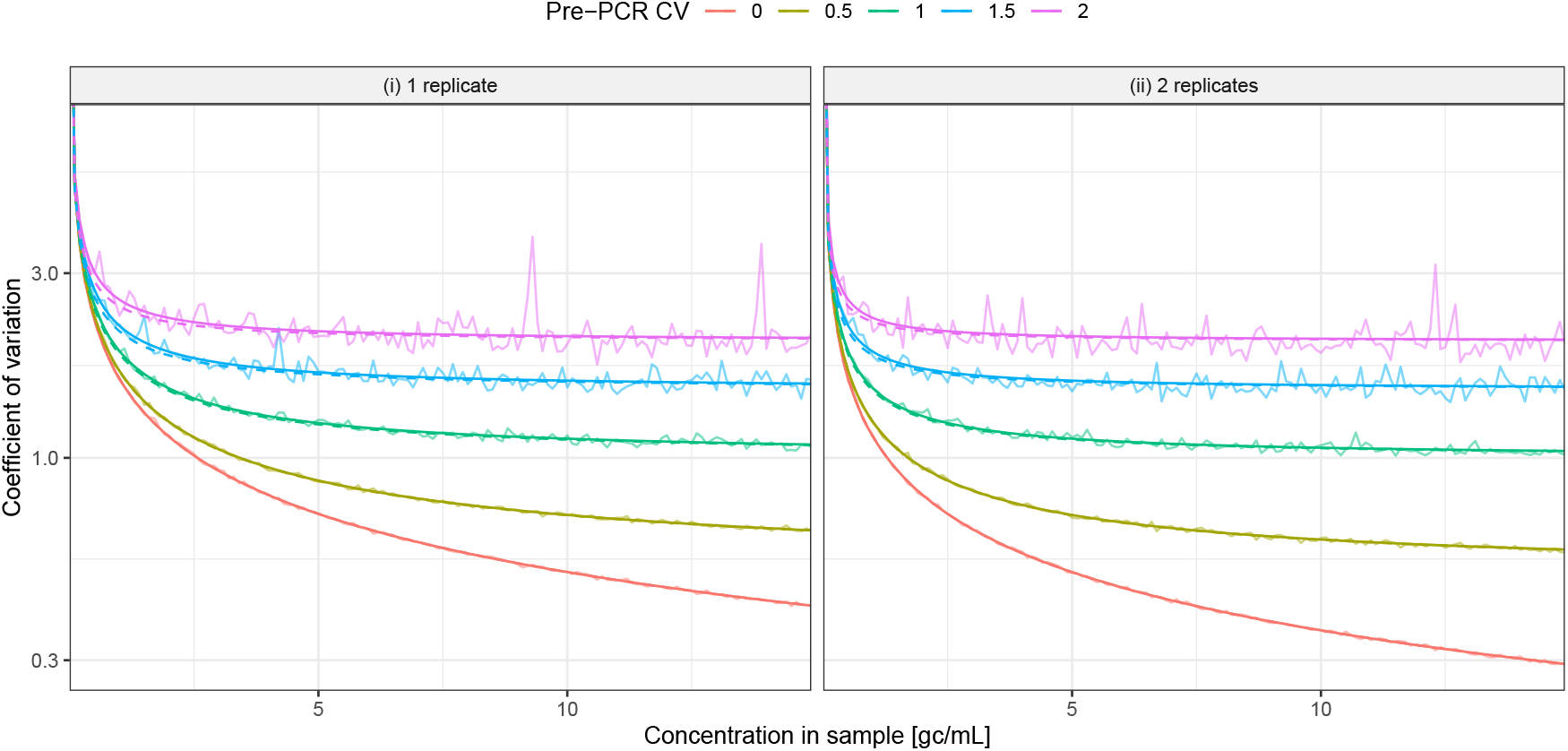
Relationship between concentration and coefficient of variation of dPCR measurements under log-normally distributed pre-PCR noise. Shown is the coefficient of variation (CV) of measurements from digital PCR for (i) a single replicate and (ii) the arithmetic mean of two replicates, as a function of the sample concentration *λ* under log-normally distributed pre-PCR noise. Jagged lines show the CV of simulated measurements for different concentrations in steps of 0.1 gc*/*mL. Straight lines show the corresponding CV as approximated using a Taylor series expansion, and dashed lines show an alternative approximation of the log-normal MGF by Asmussen et al. [3]. Colors correspond to different strengths of pre-PCR noise, as measured by the pre-PCR coefficient of variation *ν*_pre_.

Figure S6 shows a sensitivity analysis of the CV as a function of the sample concentration to the conversion factor *c* for the dPCR assay. Compared are results for a conversion factor 10x as large (green), and 10x as small (blue) as assumed in the main analysis (red). The simulated and predicted CV values are based on an assumed pre-PCR coefficient of variation of 0.5. As can be seen, a larger conversion factor reduces the CV, while a smaller conversion factor increases the CV.

Figure S7 shows a sensitivity analysis of the CV as a function of the sample concentration to the total number of droplets in the dPCR assay. Compared are results for 15000 partitions less (green), and 15000 partitions more (blue) as assumed in the main analysis (red). The simulated and predicted CV values are based on an assumed pre-PCR coefficient of variation of 0.5. As can be seen, a larger number of partitions reduces the CV, while a smaller number of partitions increases the CV.

**Fig S5.**
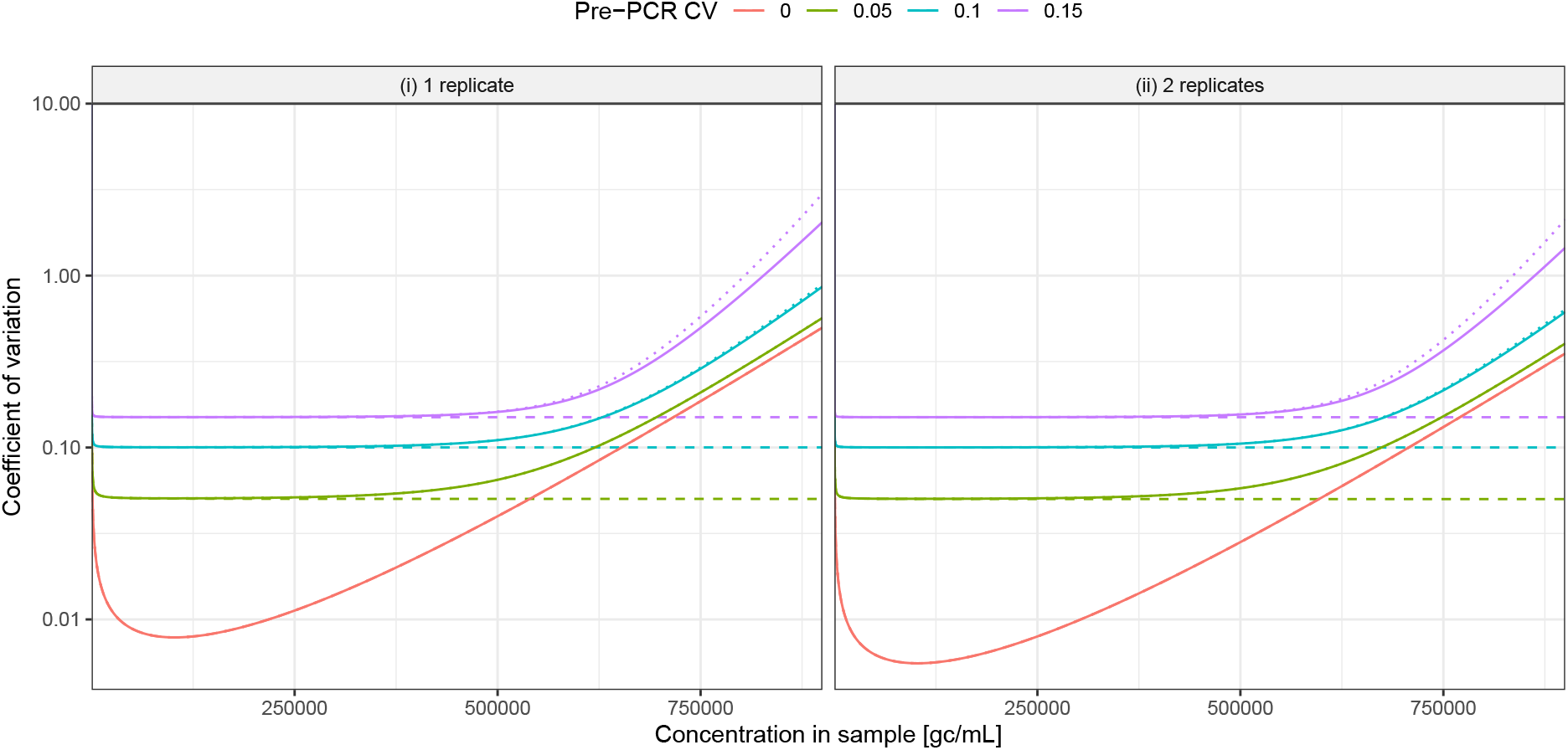
Relationship between concentration and coefficient of variation of dPCR measurements across wide range of concentrations. Shown is the coefficient of variation (CV) of measurements from digital PCR for (i) a single replicate and (ii) the arithmetic mean of two replicates, as a function of the sample concentration *λ* under gamma distributed pre-PCR noise for different concentrations. Solid lines show the CV as predicted under gamma distributed pre-PCR noise, dashed lines show an approximation using a Taylor series expansion, and dotted lines show an approximation under log-normally distributed pre-PCR noise. Colors correspond to different strengths of pre-PCR noise, as measured by the pre-PCR coefficient of variation *ν*_pre_.

**Fig S6.**
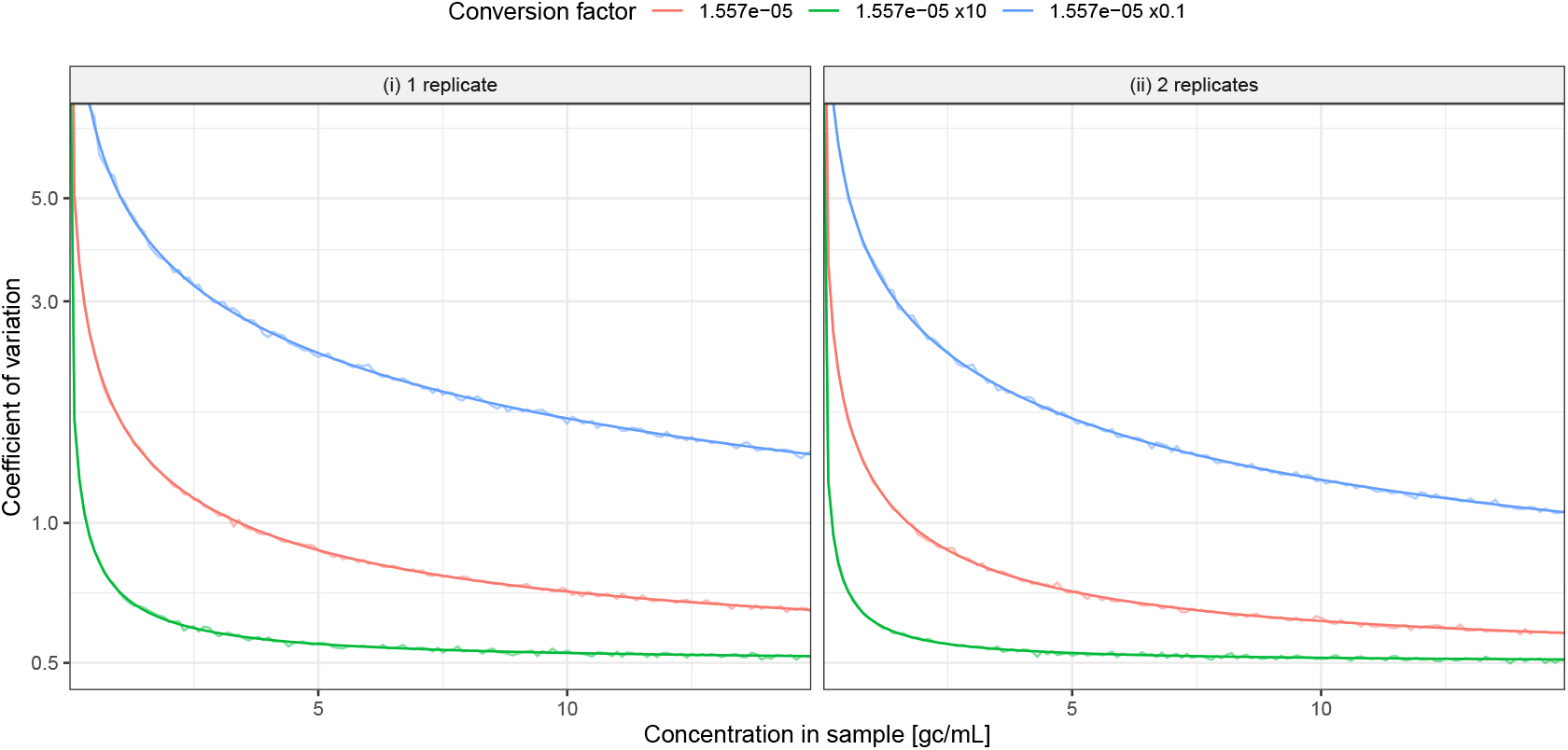
Sensitivity of the coefficient of variation of dPCR measurements to the conversion factor. Shown is the coefficient of variation (CV) of measurements from digital PCR for a conversion factor as in the main analysis (*c* = 30 ·0.519 nL, red), 10x as large (green), and 10x as small (blue). The CV is shown for (i) a single replicate and (ii) the arithmetic mean of two replicates, as a function of the sample concentration *λ* under gamma distributed pre-PCR noise with a pre-PCR coefficient of variation of 0.5. Jagged lines show the CV of simulated measurements for different concentrations in steps of 0.1 gc*/*mL, straight lines show the corresponding CV as predicted by Eq. (16).

**Fig S7.**
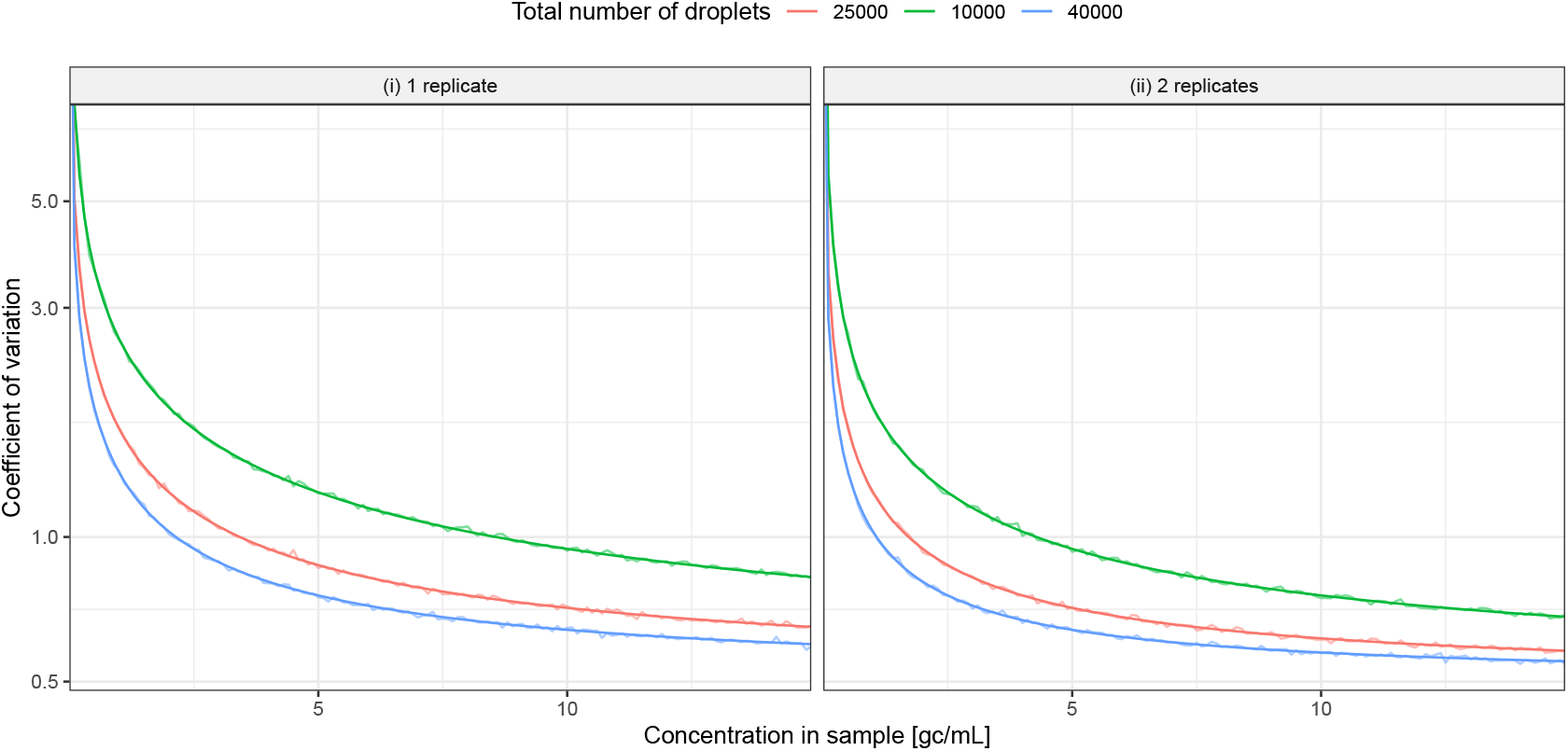
Sensitivity of the coefficient of variation of dPCR measurements to the total number of partitions. Shown is the coefficient of variation (CV) of measurements from digital PCR for 25000 total partitions, i. e. as in the main analysis (red), 10000 total partitions (green), and 40000 total partitions (blue). The CV is shown for i) a single replicate and (ii) the arithmetic mean of two replicates, as a function of the sample concentration *λ* under gamma distributed pre-PCR noise with a pre-PCR coefficient of variation of 0.5. Jagged lines show the CV of simulated measurements for different concentrations in steps of 0.1 gc*/*mL, straight lines show the corresponding CV as predicted by Eq. (16).

#### D.2 Probability of non-detection

Figure S9 shows the probability of non-detection as a function of the sample concentration under gamma distributed pre-PCR noise. The presence of pre-PCR noise considerably increases the probability of zero measurements. Moreover, from Eq. (24) we see that the probability of non-detection decreases with the number of partitions and the number of replicates.

**Fig S8.**
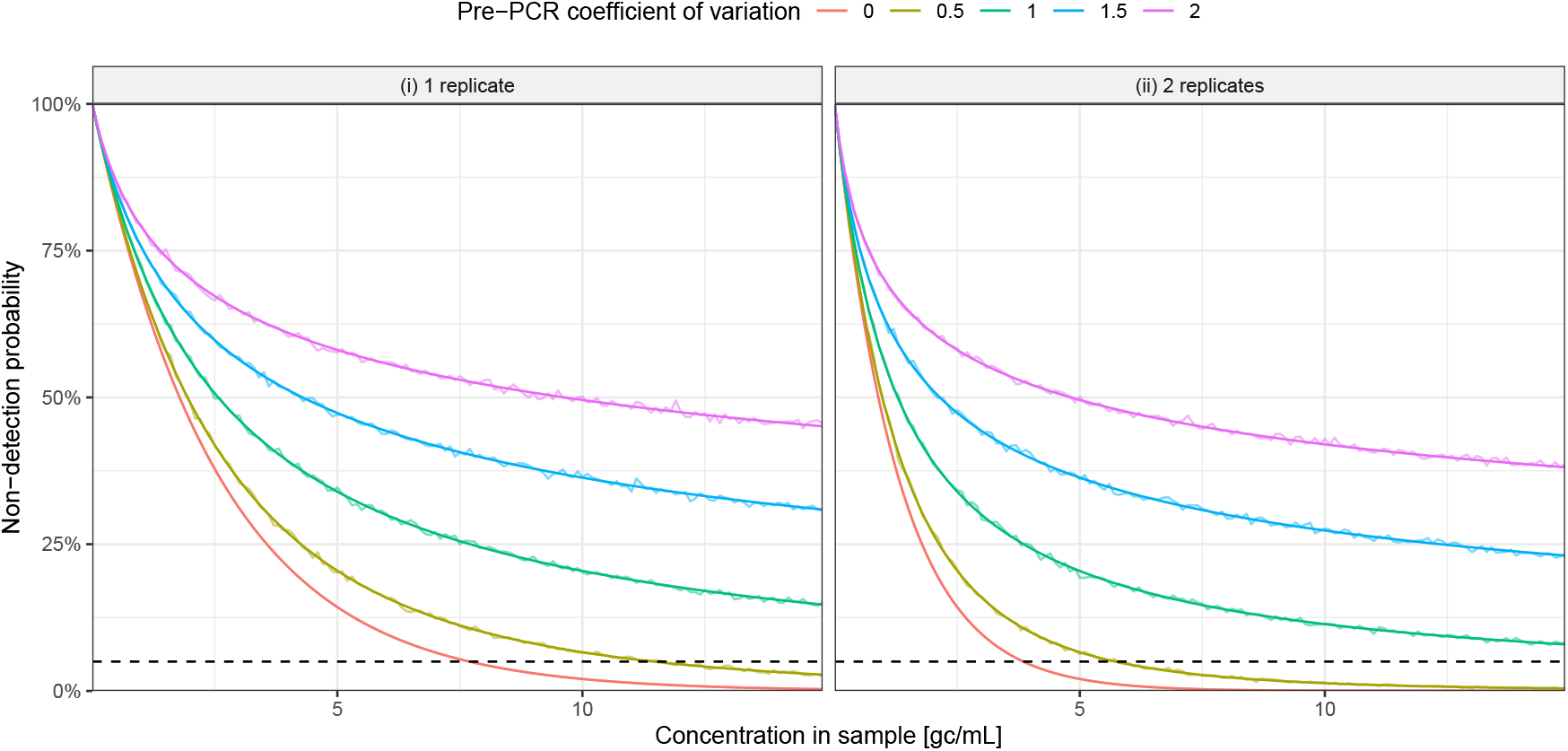
Relationship between concentration and probability of non-detection of dPCR measurements under gamma distributed pre-PCR noise. Shown is the probability of a zero measurement (i. e. non-detection) in digital PCR for (i) a single replicate and (ii) the arithmetic mean of two replicates, as a function of the sample concentration *λ* under gamma distributed pre-PCR noise. Jagged lines show the probability of non-detection of simulated measurements for different concentrations in steps of 0.1 gc*/*µL, while straight lines show the corresponding probability of non-detection as predicted by Eq. (24). Dashed horizontal lines indicate the limit of detection (LoD), defined as a 5% probability of non-detection. Colors correspond to different strengths of pre-PCR noise, as measured by the pre-PCR coefficient of variation *ν*_pre_.

Figure S9 shows the probability of non-detection as a function of the sample concentration under log-normally instead of gamma distributed pre-PCR noise. In addition to the CV of simulated measurements, a theoretical prediction using an approximation of the log-normal MGF by Asmussen et al. [3] (see Supplement C.2) is shown. As can be seen, the approximation matches the simulated CV values well, but slightly underestimates the CV at low concentrations when the pre-PCR variation is very high (Figure S9). Compared to the case of gamma distributed pre-PCR noise (Figure S8), the probability of non-detection falls considerably faster under log-normally distributed pre-PCR noise.

**Fig S9.**
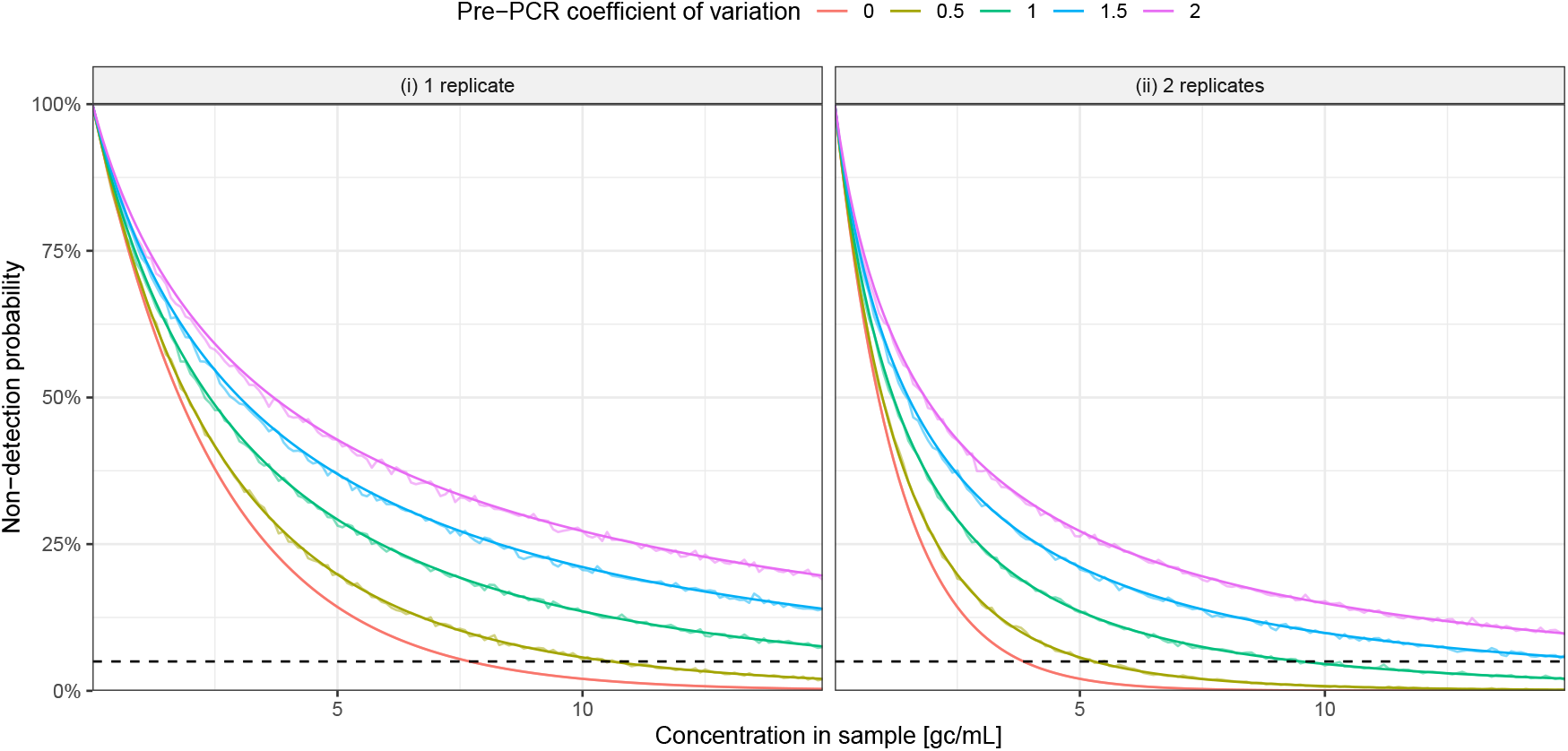
Relationship between concentration and probability of non-detection of dPCR measurements under log-normally distributed pre-PCR noise. Shown is the probability of a zero measurement (i. e. non-detection) in digital PCR for (i) a single replicate and (ii) the arithmetic mean of two replicates, as a function of the sample concentration *λ* under log-normally distributed pre-PCR noise. Jagged lines show the probability of non-detection of simulated measurements for different concentrations in steps of 0.1 gc*/*µL, while straight lines show the corresponding probability of non-detection as predicted using an approximation of the log-normal MGF by Asmussen et al. [3]. Dashed horizontal lines indicate the limit of detection (LoD), defined as a 5% probability of non-detection. Colors correspond to different strengths of pre-PCR noise, as measured by the pre-PCR coefficient of variation *ν*_pre_.

### E Comparison with empirical data

We used dPCR data of pathogen concentrations in Swiss wastewater to validate our statistical model of dPCR measurements. For this we used data from 3031 autosampler-based, 24-hour-composite samples taken at 14 different municipal treatment plants throughout Switzerland in the time period July 2023 – June 2024. For each sample, viral RNA was extracted from a 40 mL aliquot by vacuum filtration, yielding an elution of 80 µL, and subsequently diluted by a factor of 1:3. As described in Huisman et al. [6] and Nadeau et al. [7], for quantification via dPCR, a 25 µL reaction mix with 5 µL of the template and 20 µL of reagents was run in a droplet-based, sixplex dPCR assay with Influenza A (M gene), Influenza B (M gene), Respiratory Syncytial Virus (N gene), SARS-CoV-2 (N1 and N2 genes), and an internal extraction efficiency control as targets. Quantification was performed using the Naica Crystal Digital PCR System (Stilla^®^ Technologies) with a Sapphire Chip with a maximum number of 30000 droplets and an average droplet volume of 0.519 nL.

For each target and sample, the CV was empirically estimated from two technical replicates, using the formula

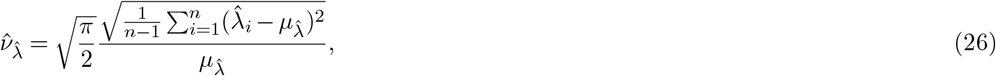

where 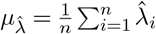 is the empirical mean of the replicates and 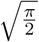 is a bias correction factor for the empirical standard deviation. We note that 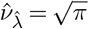 in the special case where one of the replicate measurements is zero. To estimate the probability of non-detection, we used the technical replicates by treating the first replicate as an estimate of the true target concentration, and the second replicate as an indicator of the probability of zero. Specifically, for a given window *w* ∈ {0,…, 30}, we selected all replicate pairs where the result of the first replicate was between *w* and *w* + 1 gc*/*mL. Of these, we then computed the percentage of second replicates with a result of zero to estimate *p*_zero_(*w* + 0.5). The moving window was here used to group samples with a similar concentration together while accounting for the noise of the measurements. Given the empirical estimates for different target concentrations, we applied locally estimated scatterplot smoothing (LOESS) [8] to estimate a smooth relationship of the target concentration with the CV and probability of non-detection, respectively.

Figure S10 shows the comparison between theoretical predictions and the real-world dPCR measurements for Influenza A and B virus, RSV, and SARS-CoV-2.

**Fig S10.**
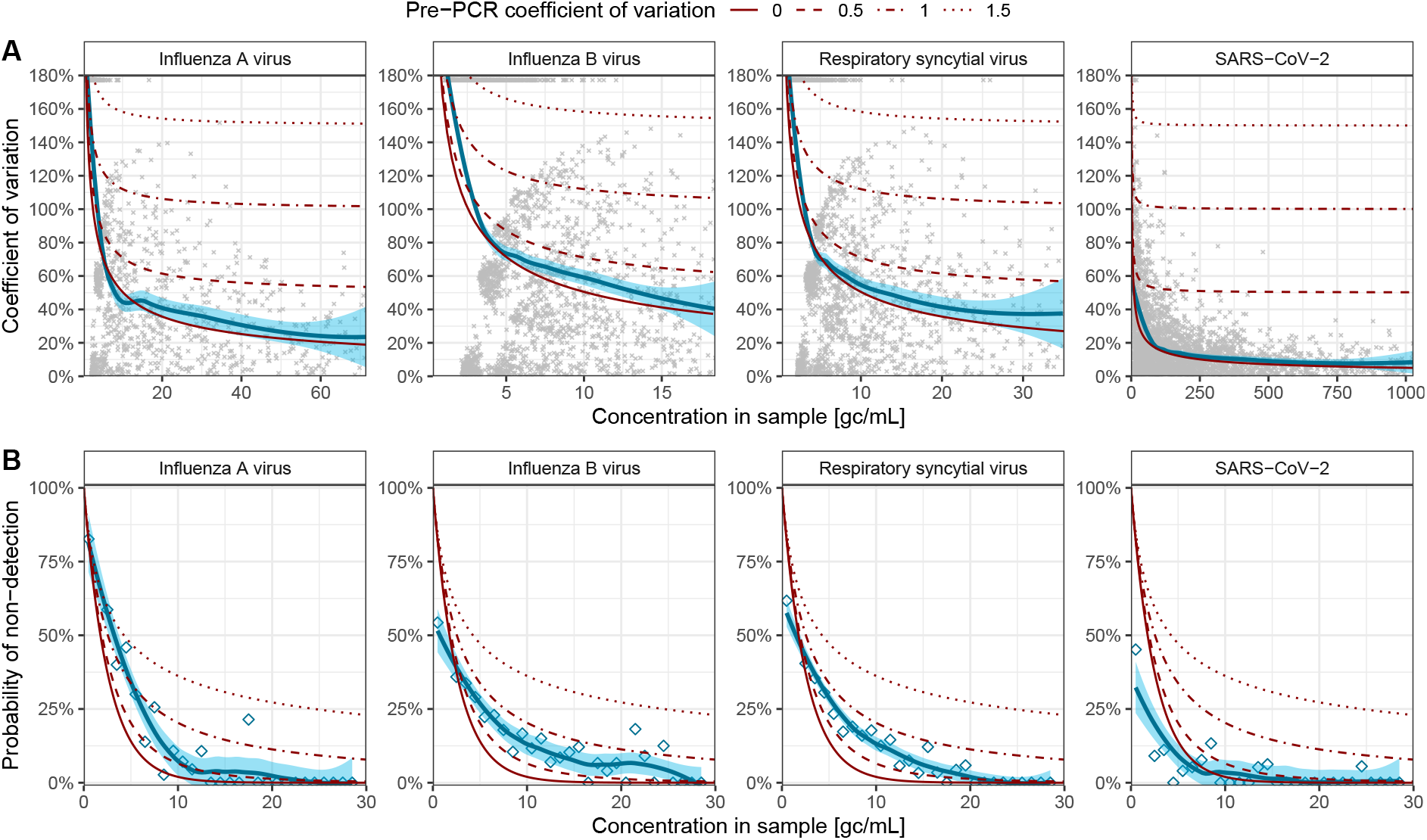
Comparison of theoretical predictions with real-world dPCR measurements. Shown are empirical estimates of (A) the coefficient of variation and (B) the probability of non-detection for dPCR measurements of Influenza A and B virus, RSV, and SARS-CoV-2 concentrations from 14 wastewater treatment plants in Switzerland. Coefficient of variation (CV) estimates are based on the bias-corrected empirical standard deviation of two technical replicates, respectively, divided by their empirical mean (dots). Probability of non-detection estimates are based on the percentage of second replicates with a result of zero among all replicate pairs where the result of the first replicate falls within a moving window of 1 gc*/*mL width (diamonds). Shown in blue with 95% uncertainty intervals are LOESS-based kernel regressions of the empirical estimates, indicating the relationship between the concentration in the sample and empirical CV and probability of non-detection, respectively. Shown in red are the CV and probability of non-detection as predicted by the dPCR-specific measurement model, for different strengths of the pre-PCR coefficient of variation *ν*_pre_ under gamma distributed pre-PCR noise (dashed and dotted lines). Note that as the empirical CV is based on technical replicates of the same sample, respectively, it does not include variation from pre-PCR noise.

### F Likelihoods for non-zero dPCR measurements

#### F.1 Conditional distribution of non-zero measurements

In the proposed hurdle model, *f*_PCR_(*x*) is the distribution of measurements conditioned on a non-zero result. While the overall distribution of measurements has mean *λ* and coefficient of variation 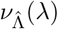, this is not true for the conditional distribution. Instead, using the law of total expectation, i. e.

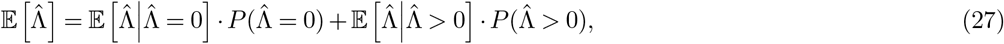

we know that

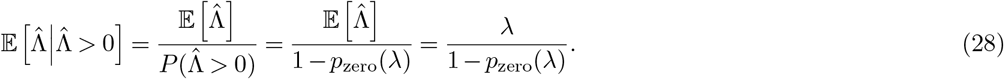

That is, the conditional mean increases relative to the unconditional mean as the probability of non-detection becomes large. Moreover, by the same argument, we see that

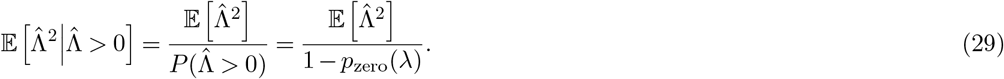

Expanding the conditional variance and inserting the above, we get

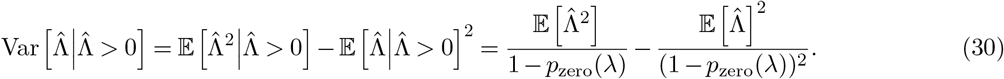

Then, using 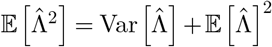, we obtain

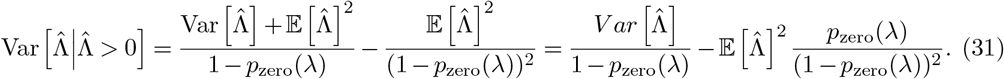

From the conditional variance, we can derive the conditional coefficient of variation as

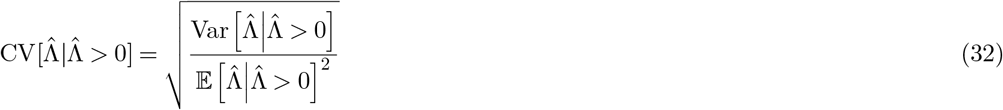

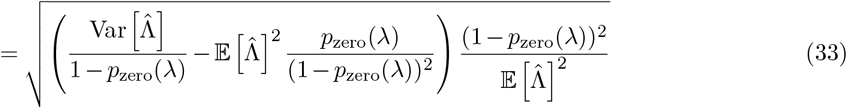

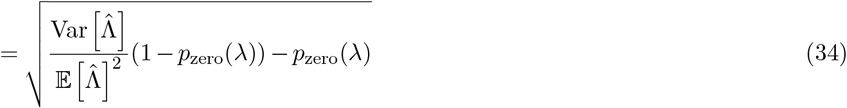

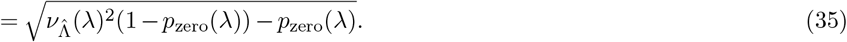

Thus, the conditional CV decreases relative to the unconditional CV as the probability of non-detection becomes large. The resulting relationship between the unconditional and conditional mean and CV is also visualized in Figure S11.

**Fig S11.**
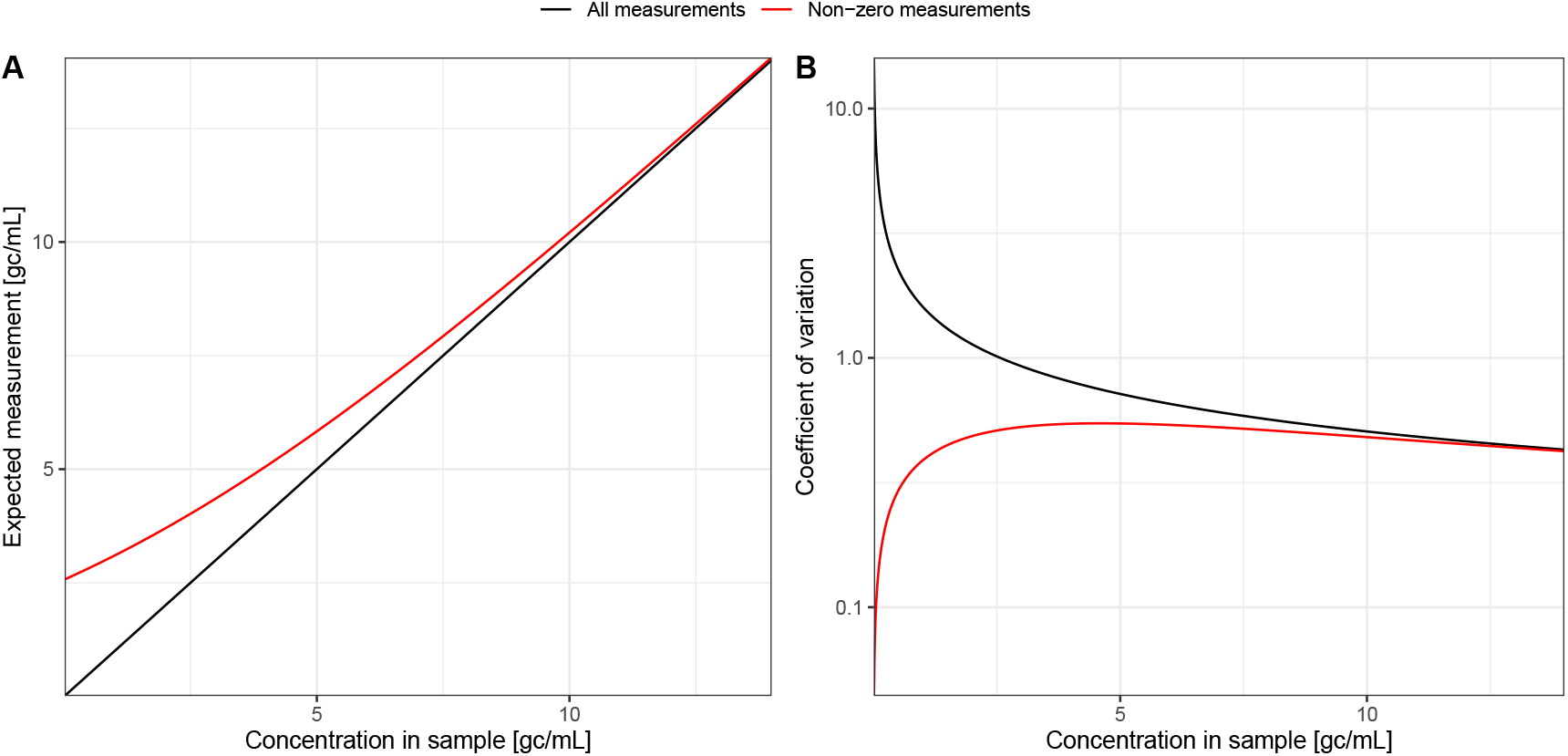
Effect of conditioning on non-zero measurements on the mean and coefficient of variation of measurements. Shown are the (A) mean and (B) coefficient of variation of the unconditional (black, all measurements) and conditional (red, non-zero measurements) distribution of measurements as a function of the true concentration in the sample. The differences between the unconditional and conditional distribution increase as the concentration becomes small.

#### F.2 Continuous vs. discrete measurement distributions

In theory, if the total number of partitions, the partition volume, and the scaling factor of each measurement are known exactly, the number of positive partitions in the PCR could be back-calculated from the concentration estimate 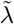 as

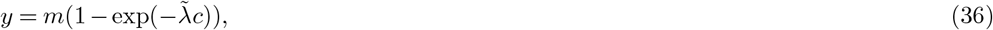

although we note that in such a situation *y* is likely already known. Given these quantities, it would also be possible to fit a Binomial distribution to the observed *y* according to Eq (3) from the main text. The goal of this work was however to propose a likelihood model that also works well if the specific parameters of the PCR are not exactly known. In such a situation, the use of a Binomial likelihood has two important drawbacks. First, and most importantly, when using a Binomial distribution, any misspecification or uncertainty about the PCR parameters directly influences the expected concentration estimate, not only its variance. Thus, a lack of knowledge about the PCR parameters can bias concentration estimates even if the measurements 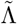 are unbiased. Second, in contrast to our continuous distribution approach, if only 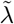 is known exactly, then the back-computation of *y* depends on the assumed PCR parameters *m, s*, and *v*, which makes a joint estimation of these parameters from dPCR data difficult.

#### F.3 Comparison of continuous measurement distributions

When using a continuous distribution to model dPCR measurements, we only approximate the true distribution of measurements, which are derived from positive partition counts and therefore discretely distributed. Figure S12 compares the cumulative distribution functions (CDFs) of different continuous probability distributions to the discrete CDF of the simulated true distribution of measurements. As can be seen, the normal distribution is an inappropriate approximation as it assigns non-zero probability to negative measurements. Since zero measurements are represented by our hurdle model, we can use strictly positive distribution to represent non-zero measurements (see Figure S12A). As strictly positive distributions, we compared the log-normal, inverse Gaussian, and gamma distributions. While the log-normal distribution is often used as a default to model positive measurements, we note that the concentration estimates 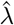 are a log-linear transform of the Binomially (and therefore approximately normally) distributed partition counts. The log-normal distribution, however, would fit well for an exponential transform. Indeed, we find that a gamma distribution offers a better fit than the log-normal or inverse Gaussian distribution, especially at larger concentrations (see Figure S12B), as indicated by their discretized Kullback-Leibler divergence to true distribution (KL_Gamma_ = 0.01, KL_Log-normal_ = 0.05, KL_Inverse Gaussian_ = 0.06). Other strictly positive distributions such as the Truncated normal distribution could also provide a suitable approximation, but are difficult to parameterize by their mean and coefficient of variation. We therefore use a gamma distribution to model non-zero measurements in this work.

**Fig S12.**
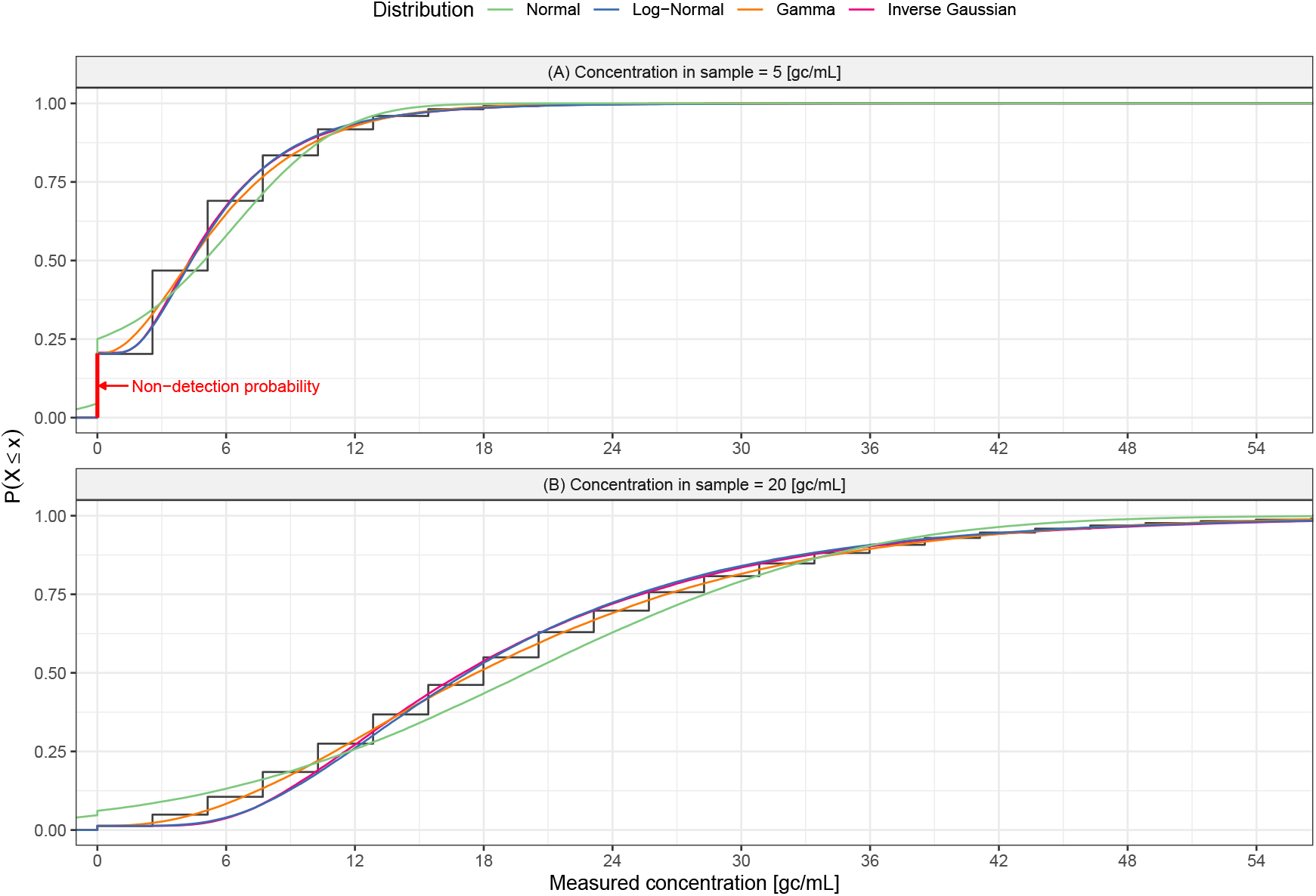
Continuous distributions approximating the distribution of dPCR measurements. Shown in black is the discrete cumulative distribution function of dPCR measurements obtained from the number of positive partitions in the assay for a concentration of (A) 5 gc*/*mL and (B) 20 gc*/*mL. This distribution can be approximated using a hurdle model with a probability of non-detection as defined in Eq. (23) and continuously distributed non-negative measurements with conditional mean 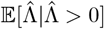 and conditional coefficient of variation 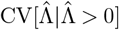. Shown in colors are the continuous cumulative distribution functions of a hurdle model using a normal (green), log-normal (blue), gamma (orange), and inverse Gaussian (pink) distribution for the non-negative measurements, respectively.

### G Inference from dPCR measurements

#### G.1 Estimation

The dPCR-specific likelihood was implemented as a function in the probabilistic programming language stan [9]. We integrated this likelihood function in a validation model for estimating a single concentration from replicate measurements, ia generalized linear model (GLM) for estimating the association of covariates with the target concentration, and in the EpiSewer wastewater model for estimating reproduction numbers and other transmission parameters from wastewater measurements over time.

We fitted models via Markov chain Monte Carlo (MCMC) using cmdstan version 2.34.1 via cmdstanr version 0.8.1 [10]. We ran four No-U-Turn sampler chains with 1,000 warm-up and 1,000 sampling iterations each, with a maximum tree depth of 15, an initial step size of 0.01, and an adaptation target acceptance statistic of 0.99. We checked the fitted models for low effective sample sizes (ESS < 400) [11], and for convergence problems using the number of divergent transitions [12], the Bayesian fraction of missing information (E-BFMI < 0.2) [13] and the Gelman-Rubin convergence diagnostic (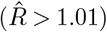) [14]. The checks indicated sufficient effective sampling sizes and good convergence and mixing of chains.

#### G.2 Validation

We validated inference via the dPCR-specific likelihood by fitting the model for a single concentration to simulated dPCR measurements with different true sample concentrations (1, 3, and 5 gc*/*mL). For comparison purposes, we also tested the model with two simpler likelihoods, i. e. i) using a normal distribution with constant variance and ii) using a log-normal distribution with constant CV. When using the log-normal likelihood, zero measurements had to be dropped from the analysis, otherwise, the models and data used were identical for all likelihoods. We deliberately used a large number of 100 simulated replicates per concentration for estimation to study asymptotic differences between the different likelihoods.

Figure S13 shows the true concentration and estimated concentrations using the dPCR-specific as well as the normal and log-normal likelihood function, together with the posterior predictive distributions for the dPCR measurements. When using a dPCR-specific likelihood, the probability of zero measurements is explicitly represented and the true concentration is estimated without bias. When using a normal likelihood, estimates of the concentration are unbiased, however, the model can predict negative measurements. When using a log-normal likelihood, no negative measurements are predicted but zero measurements must be dropped from the analysis. This leads to a significant bias in the estimated concentration when the true concentration is small.

**Fig S13.**
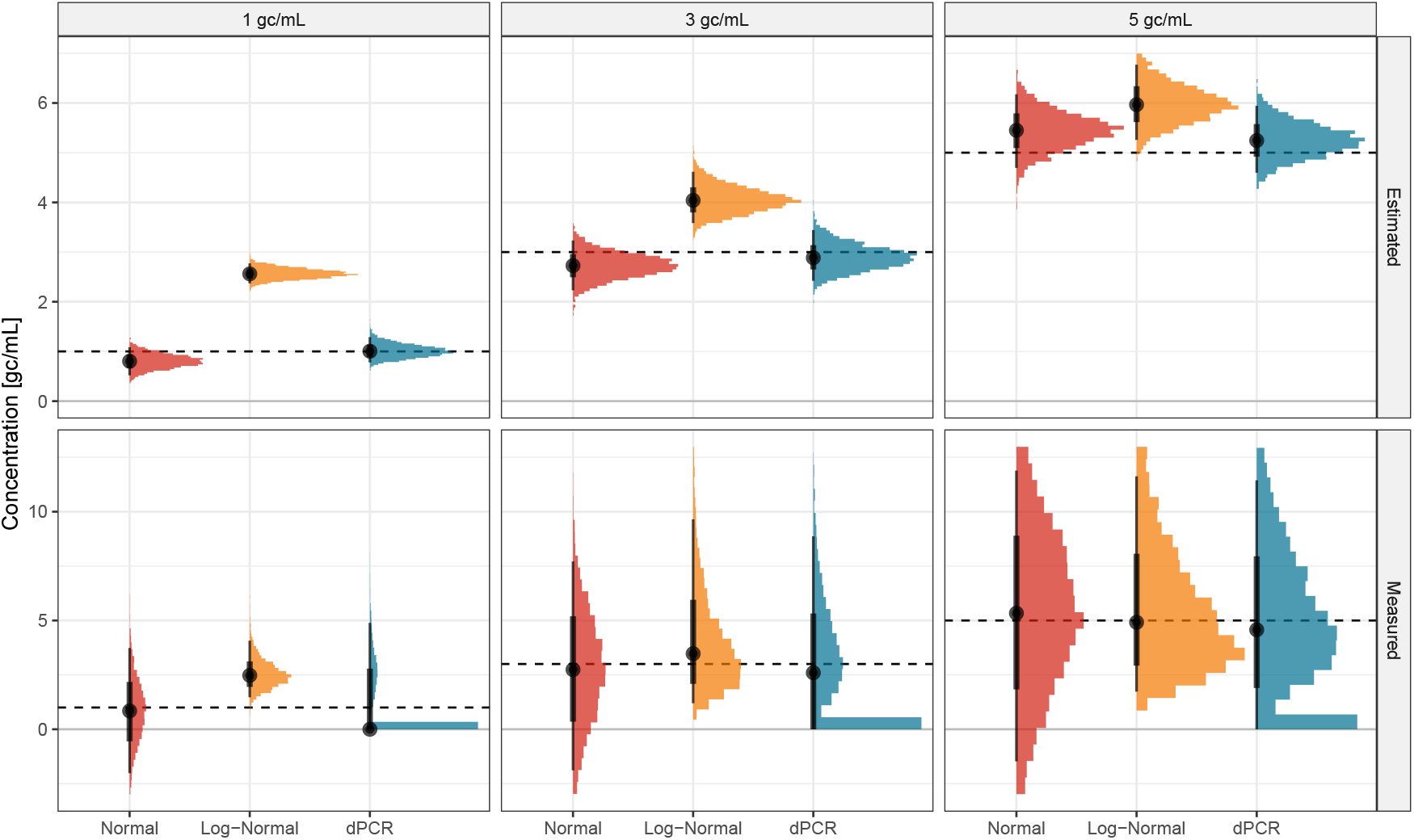
Comparison of estimated concentration and predicted dPCR measurements under different likelihood functions. Shown is the posterior distribution of the estimated true concentration (top) and the posterior predictive distribution of measurements (bottom) from a Bayesian model fitted with a normal (red), log-normal (orange), and dPCR-specific (blue) likelihood to simulated dPCR measurements of different concentrations (1, 3, and 5 gc*/*mL). Dashed lines show the true concentration, respectively.

### H Application to eDNA-based biomonitoring

We modeled the expected concentration of free-eDNA of Bugula neritina *t* hours after removal of the organism as

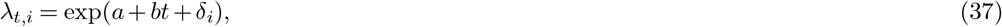

where exp(*a*) is the concentration at *t* = 0 (intercept), *b* the exponential decay factor, and exp(*δ*_*i*_) a multiplicative fixed effect for the sample location in the aquarium (back left, back right, front left, front right, and middle). The half-life period can be computed as 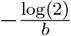. When analyzing data from all aquariums combined, we accounted for differences between aquariums using

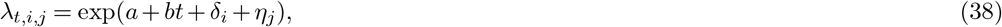

where exp(*η*_*j*_) is a multiplicative effect for the aquarium of type *j* (low, medium, or high biomass). We fitted the model with a normal, log-normal, and dPCR-specific likelihood respectively (see Supplement G.1 for estimation details). For the log-normal likelihood, we placed a half-normal prior with a standard deviation of 0.5 on the coefficient of variation. For the normal likelihood, we assumed a fixed variance, but parameterized it in terms of the average coefficient of variation to use the same noise prior as in the log-normal model. For the dPCR-specific likelihood, based on the lab protocol in Scriver et al. [15], we assumed a scaling factor of 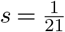 for the reaction mix and *m* = 20000 partitions with a volume of *v* = 1 nL for the BioRad200™ droplet generator without droplet loss. Again, we used a half-normal prior with a standard deviation of 0.5 for the remaining pre-PCR variation (*ν*_pre_).

To reproduce the original approach by Scriver et al., we simply fitted a normal likelihood to the averages of measurements at each point in time. As in Scriver et al., measurements above the maximum value of the negative controls in the experiment (0.08 gc*/*µL) were removed before averaging.

### I Application to wastewater-based epidemiology

To demonstrate inference from a time series of measurements, we used samples taken on average 5 times per week at the treatment plants of Zurich and Geneva, Switzerland in the time period Oct 19, 2022 – April 30, 2023, extracted under the same protocol as described in Supplement E and quantified using a similar, fourplex dPCR assay with Influenza A (M gene), Influenza B (M gene), Respiratory Syncytial Virus (N gene), and SARS-CoV-2 (N2 gene) as targets. For each treatment plant separately, we fitted the wastewater model implemented in the EpiSewer R package to the longitudinal measurements of Influenza A virus. In this model, the effective reproduction number was smoothed using a random walk prior, and infections were assumed to arise from a stochastic renewal process with Poisson distributed noise. The full model specification is provided as reproducible code at https://github.com/adrian-lison/dPCR-observation-model-study.

#### I.1 PCR parameters

##### I.1.1 Prior for number of partitions

To specify a prior for the number of partitions *m* in the assay, we took into account that *m* varies between PCR runs due to random loss of partitions. As can be seen in Figure S14, the number of lost partitions per PCR run can be approximately modeled as log-normally distributed. In principle, *m* could thus be modeled using a hyperprior for the maximum number of partitions, and hyperpriors for the mean and standard deviation of the number of lost partitions. To simplify estimation, however, we wanted to parameterize *m* instead via hyperpriors for the mean number of partitions *µ*_*m*_ and coefficient of variation of partitions *ν*_*m*_. We thus assumed that the maximum number of partitions corresponds to *µ*_*m*_ + 3*µ*_*m*_*ν*_*m*_ and chose the parameters of the log-normal distribution for lost partitions accordingly, i. e. such that the mean number of lost partitions is 3*µ*_*m*_*ν*_*m*_ and the standard deviation is *µ*_*m*_*ν*_*m*_. As a hyperprior for *µ*_*m*_, we used Normal^+^(*µ* = 20000, *σ* = 5000) to accommodate dPCR systems with between 10000 and 30000 partitions. As a hyperprior for *ν*_*m*_, we used Normal^+^(*µ* = 0, *σ* = 0.05), reflecting our expectation of less than 10% variation in the number of partitions.

**Fig S14.**
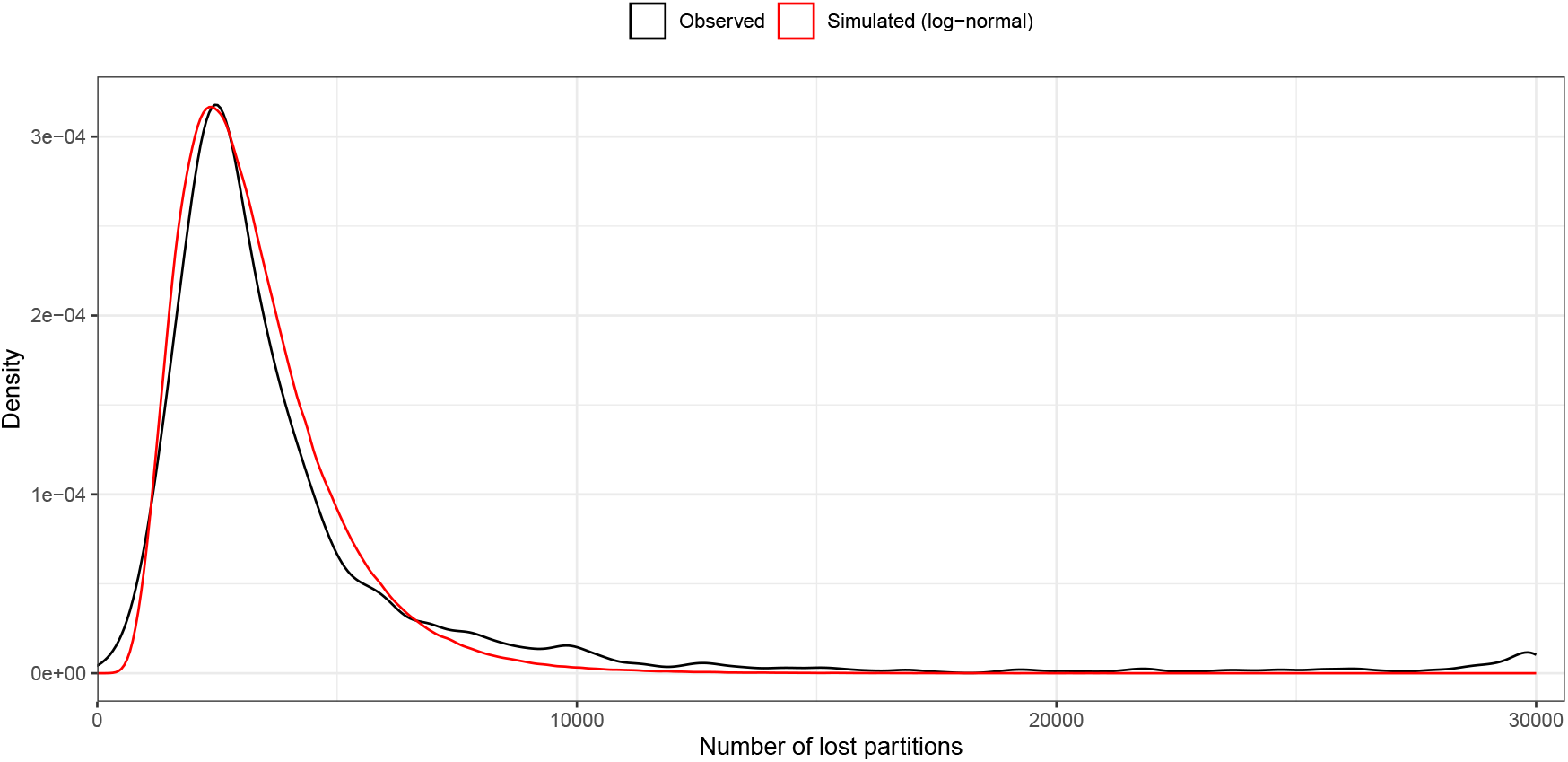
Distribution of lost partitions in a dPCR assay. Shown in black is a kernel density estimate for the distribution of the number of lost partitions in a dPCR assay based on 2733 runs on a Stilla^®^ Sapphire chip with a maximum of 30000 partitions. Shown in red is a log-normal distribution matched to the observed density.

##### I.1.2 Prior for conversion factor

Since the scaling factor *s* and partition volume *v* are not individually unidentifiable, we specified a prior for the “conversion factor” factor *c* = *sv*. This factor expresses how many expected gene copies per partition a concentration of 1 gc*/*mL in the original sample corresponds to. Here, we used a broad prior for *c* for the case of digital droplet PCR based on a typical laboratory protocol for wastewater-based epidemiology. Specifically, we assumed that extraction volumes from wastewater samples are a few dozen mL (e. g. 40mL in the assay described in Supplement E). From these, an elution of several dozen uL is produced, which corresponds to an increase in concentration by up to a factor of 1000. Due to dilution and adding of reagents for the PCR reaction, this concentration is however decreased, typically by at least a factor of 10 (e. g. by a factor of 15 in the assay described in Supplement E). This means that overall the scaling *s* in concentration through preprocessing is likely less than by a factor of 100. Importantly, while there may be further factors influencing the relationship between the original concentration in the sample and the concentration in the PCR, e. g. recovery efficiency or inhibition, *s* should only include factors that were accounted for by the lab when reporting the sample concentration. Moreover, the droplet volume *v* in digital droplet PCR is typically in the nanoliter range. Overall, we therefore expect *c* to be below 100 × 10^−6^ = 1 × 10^−4^, but typically much lower than that. For example, in the assay described in Supplement E, the conversion factor is 0.173 × 10^−4^. Based on these assumptions, we used a Normal^+^ *µ* = 1 × 10^−5^, *σ* = 4 × 10^−5^ prior for *c*, which gives a rather flat prior that has most of its probability mass below 1 × 10^−4^.

##### I.1.3 Prior for pre-PCR coefficient of variation

As a prior for *ν*_pre_, we used a standard regularizing prior, i. e. *ν*_pre_ ∼ Normal^+^(0, 1).

##### I.1.4 Estimated parameters

Figure S15 shows prior and posterior distributions for the average number of partitions, the conversion factor, and the pre-PCR coefficient of variation based on a model fitted to real-world measurements of Influenza A virus as presented in the main text.

#### 1.2 Forecasts

One week-ahead forecasts in EpiSewer were produced by assuming a continuation of the recent transmission dynamics. For this, the random walk model estimated for *R*_*t*_ was projected forward, which assumes that *R*_*t*_ will remain at its current value in expectation, but could vary from this depending on the estimated random walk variance. Thereby, the uncertainty of the projected *R*_*t*_ is higher for forecasts further into the future. Given the projected *R*_*t*_ estimates, all other latent variables such as the number of infections and the expected load in wastewater could be sampled from the generative model of EpiSewer. Since we conducted retrospective forecasts, we could use the observed flow volumes at the respective treatment plants also for the forecast dates to compute the expected concentration from on the predicted load. Finally, the sampling model corresponding to the respective likelihood function was used to predict dPCR measurements based on the forecast. This procedure was repeated for each posterior sample of the fitted model, yielding a posterior predictive distribution of *R*_*t*_, loads, and measurements.

**Fig S15.**
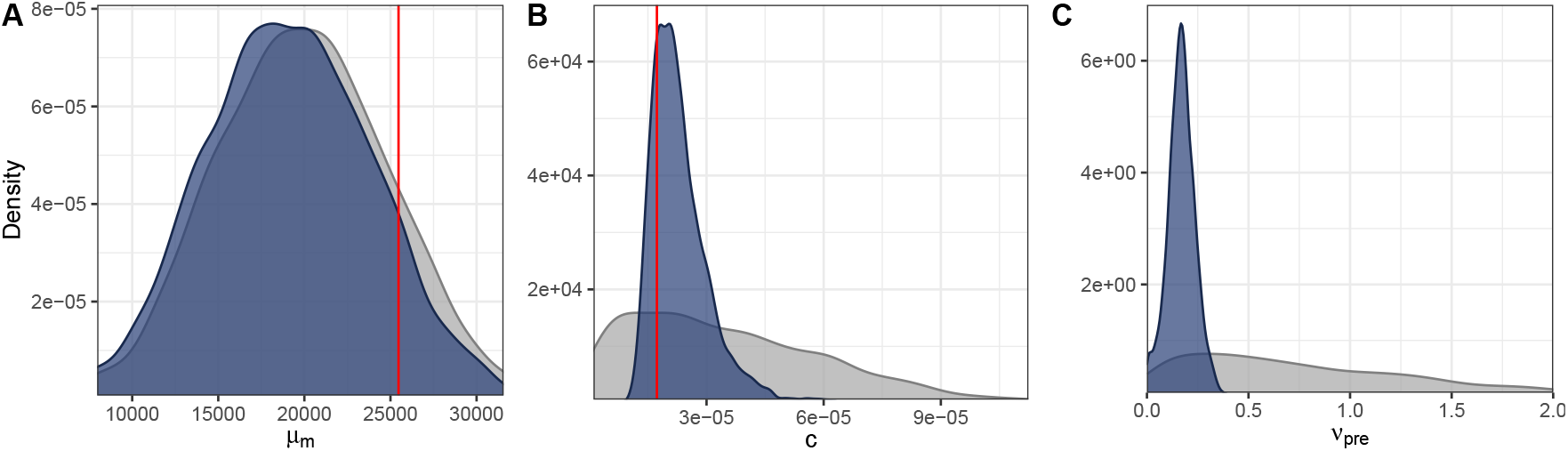
Inference of dPCR parameters and pre-PCR variation from a time series of dPCR measurements: Shown are prior and posterior distributions for (A) the average number of partitions *µ*_*m*_, the conversion factor *c*, and (C) the pre-PCR coefficient of variation *ν*_pre_ of a model applied to dPCR measurements of Influenza A virus concentrations at the municipal treatment plant of Zurich, Switzerland (Oct 19 – Dec 12, 2022). Vertical dashed lines show the true *µ*_*m*_ and *c* values based on the laboratory setup.

We assessed the forecasts of concentration measurements using the Continuous ranked probability score (CRPS) [16], i. e.

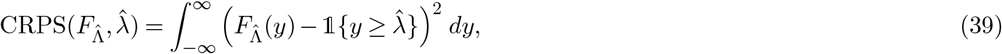

where 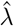 is the observed dPCR measurement and 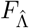 is the cumulative distribution function of the corresponding probabilistic forecast by EpiSewer. To compute the CRPS from posterior samples of our model fitted with MCMC, we used a method by Jordan et al. [17] implemented in the R package “scoringutils” [18]. We summarized the CRPS scores from different days using the arithmetic mean, denoted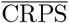.

#### I.3 Additional results

In the following, we provide additional results from fitting models using 1) a normal likelihood with constant variance, 2) a log-normal likelihood with constant coefficient of variation, and 3) the dPCR-specific likelihood on different date ranges of measurements at the treatment plants of Zurich and Geneva, Switzerland (Figures S16–S20). We also show a comparison of estimates obtained using the dPCR-specific likelihood with and without exact knowledge of the dPCR parameters (Figure S21).

**Fig S16.**
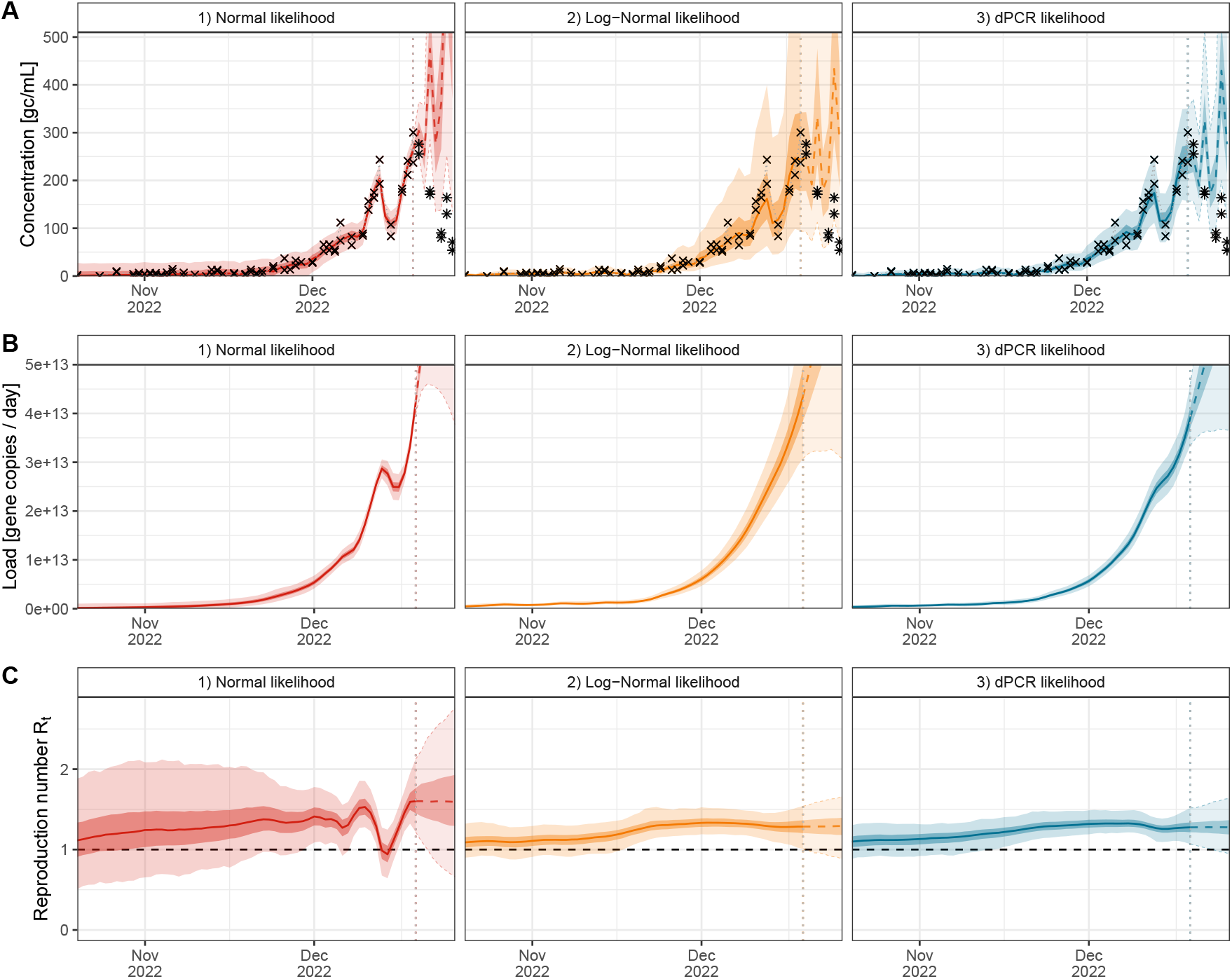
Comparison of likelihood functions on dPCR measurements of Influenza A virus concentrations at the municipal treatment plant of Zurich, Switzerland, Oct 20 – Dec 19, 2022: Shown are the results of fitting an epidemiological wastewater model (EpiSewer) to duplicate dPCR measurements of Influenza A virus at the municipal treatment plant of Zurich, Switzerland (Oct 20 – Dec 19, 2022) using either 1) a normal likelihood (red), 2) a log-normal likelihood (orange), and 3) a dPCR-specific likelihood (blue) for the observations. Each panel shows the median (lines), and 50% and 95% credible intervals (strong and weakly shaded areas) of (A) posterior predictive distributions for dPCR measurements, (B) estimated viral loads in wastewater over time, and (C) the estimated effective reproduction number over time. Estimates shown beyond Dec 26, 2022 (vertical dotted line) are forecasted based on a continuation of the latest transmission dynamics. Panel A also shows the true observed (crosses) and future (stars) dPCR measurements.

**Fig S17.**
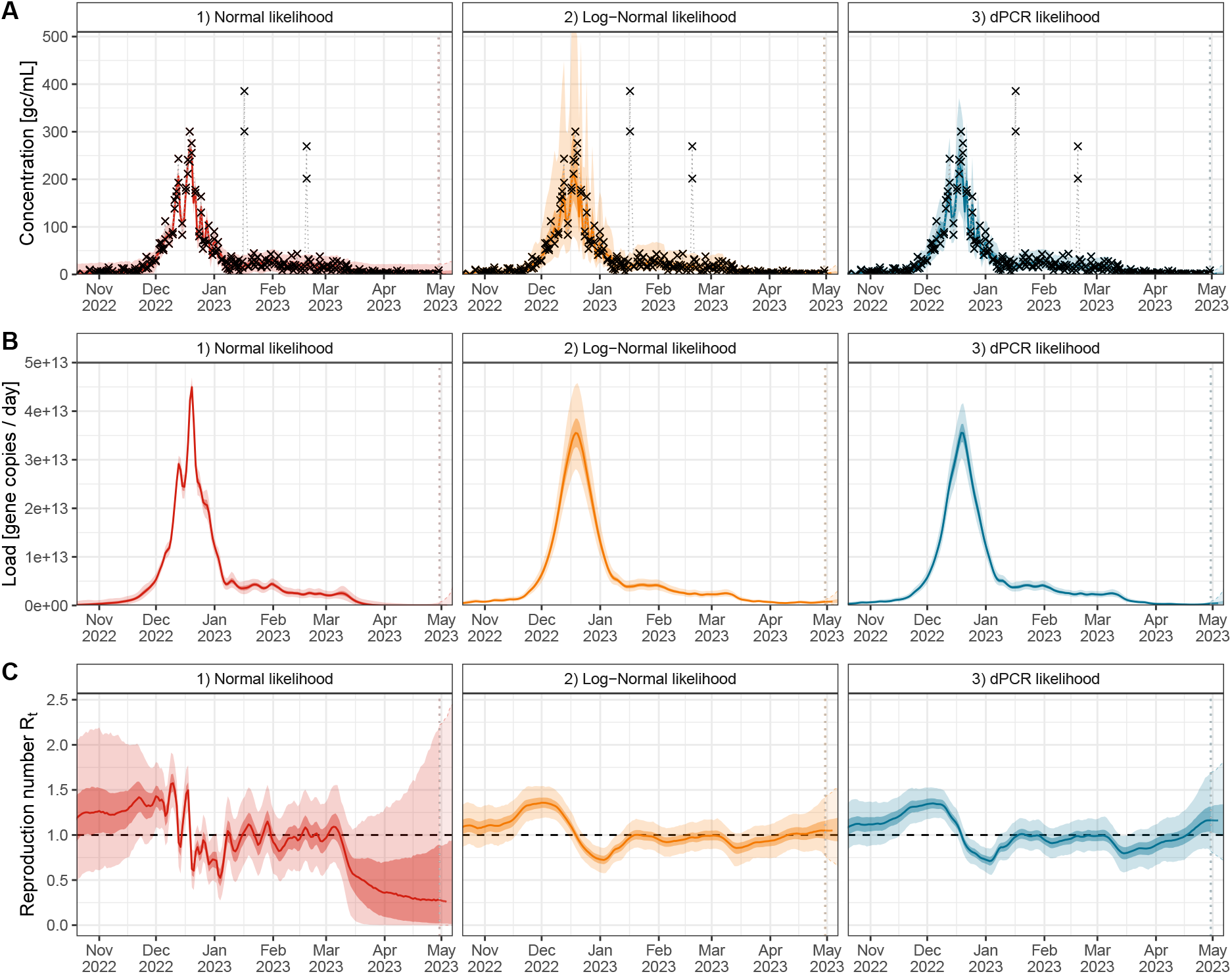
Comparison of likelihood functions on measurements of Influenza A virus concentrations at the municipal treatment plant of Zurich, Switzerland, Oct 20 – May 01, 2023: Shown are the results of fitting an epidemiological wastewater model (EpiSewer) to duplicate concentration measurements of Influenza A virus at the municipal treatment plant of Zurich, Switzerland (Oct 20 – May 01, 2023) using either 1) a normal likelihood (red), 2) a log-normal likelihood (orange), and 3) a dPCR-specific likelihood (blue) for observed measurements. Each panel shows the median (lines), and 50% and 95% credible intervals (strong and weakly shaded areas) of (A) posterior predictive distributions for concentration measurements, (B) estimated viral loads in wastewater over time, and (C) the estimated effective reproduction number over time. Estimates shown beyond Dec 26, 2022 (vertical dotted line) are forecasted based on a continuation of the latest transmission dynamics. Panel A also shows the true observed (crosses) and future (stars) measurements, respectively. Measurements on January 17, 2023, and February 2, 2023 were removed as outliers before model fitting.

**Fig S18.**
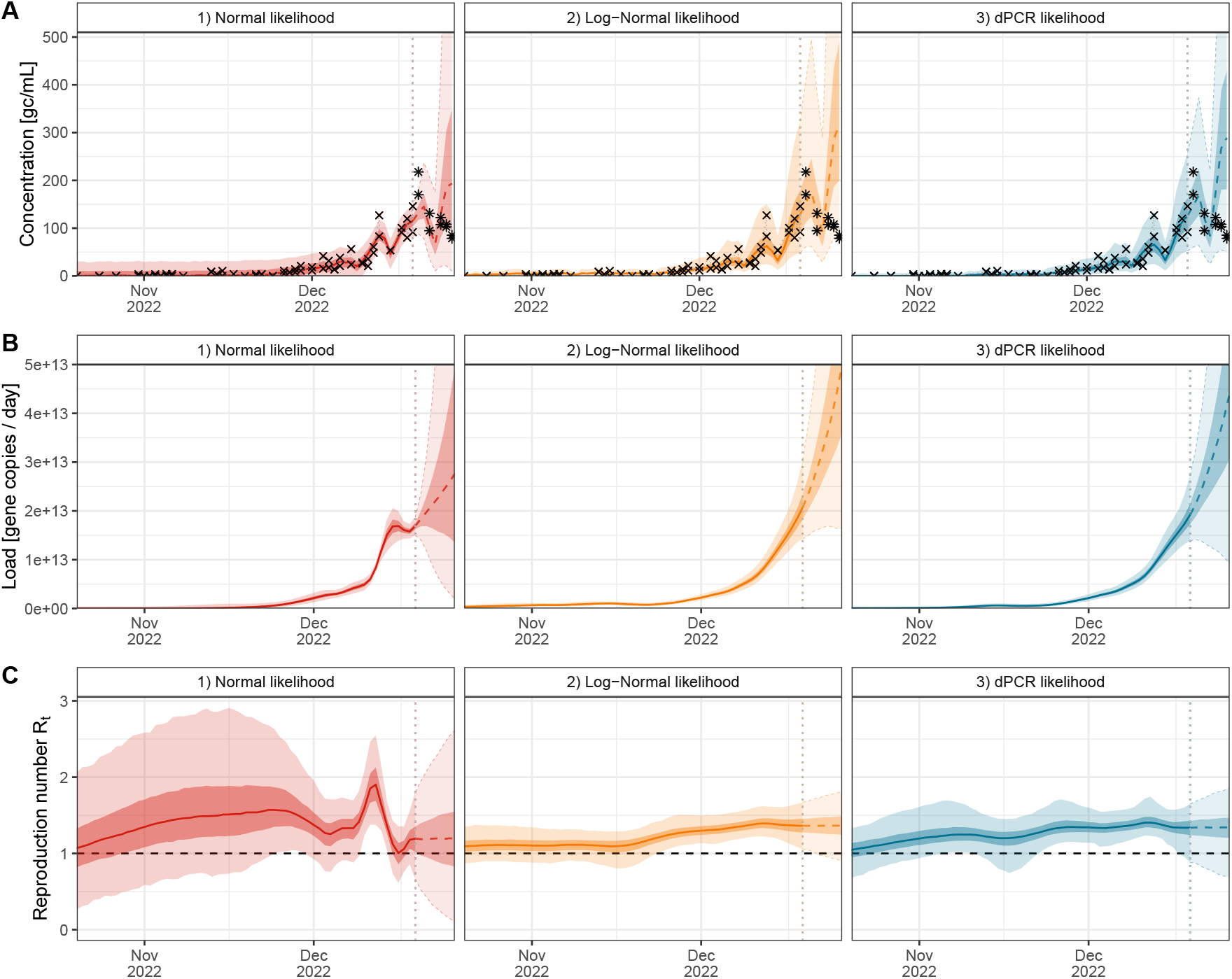
Comparison of likelihood functions on measurements of Influenza A virus concentrations at the municipal treatment plant of Geneva, Switzerland, Oct 20 – Dec 19, 2022: Shown are the results of fitting an epidemiological wastewater model (EpiSewer) to duplicate dPCR measurements of Influenza A virus at the municipal treatment plant of Geneva, Switzerland (Oct 20 – Dec 19, 2022) using either 1) a normal likelihood (red), 2) a log-normal likelihood (orange), and 3) a dPCR-specific likelihood (blue) for the observations. Each panel shows the median (lines), and 50% and 95% credible intervals (strong and weakly shaded areas) of (A) posterior predictive distributions for dPCR measurements, (B) estimated viral loads in wastewater over time, and (C) the estimated effective reproduction number over time. Estimates shown beyond Dec 26, 2022 (vertical dotted line) are forecasted based on a continuation of the latest transmission dynamics. Panel A also shows the true observed (crosses) and future (stars) dPCR measurements.

**Fig S19.**
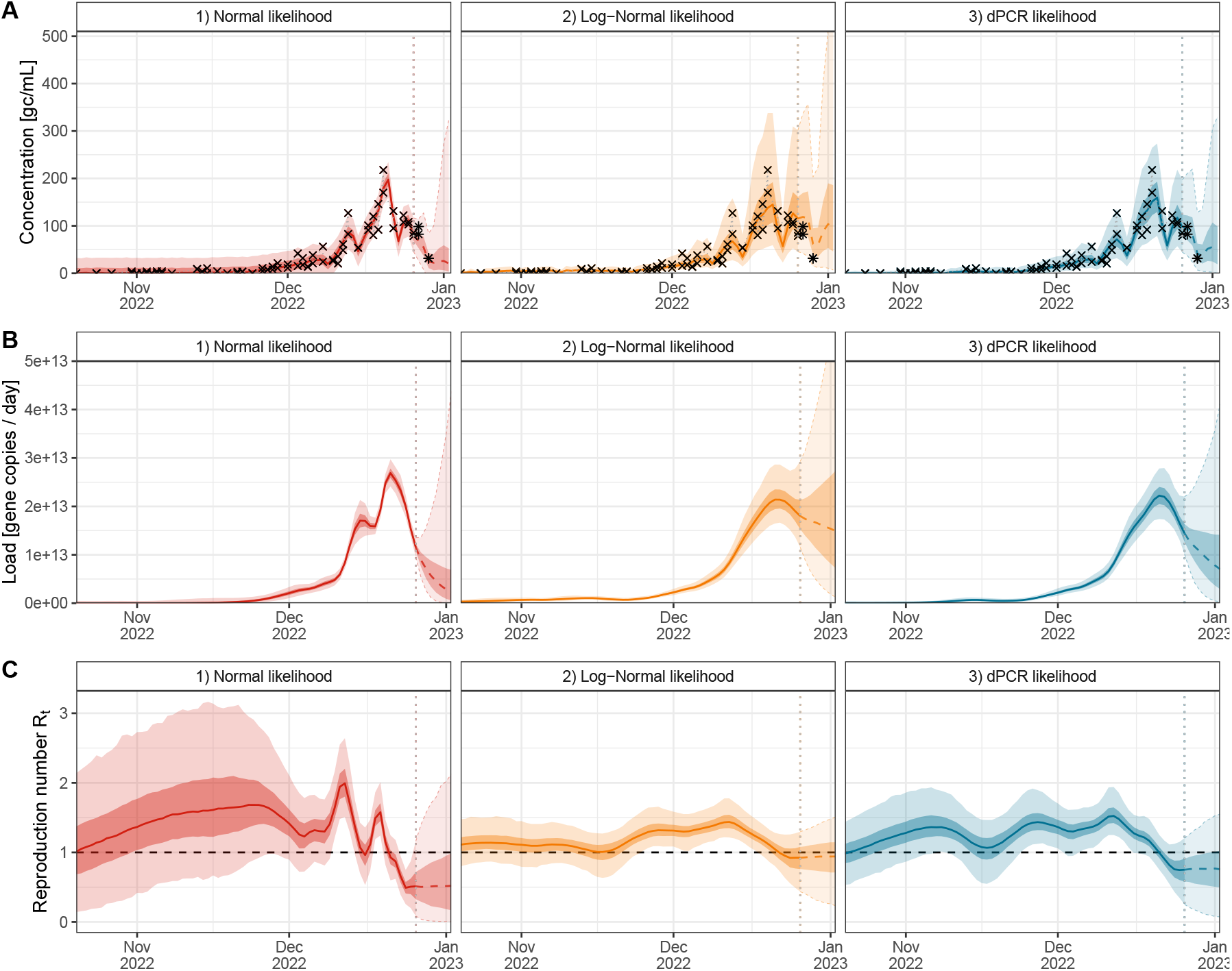
Comparison of likelihood functions on measurements of Influenza A virus concentrations at the municipal treatment plant of Geneva, Switzerland, Oct 20 – Dec 26, 2022: Shown are the results of fitting an epidemiological wastewater model (EpiSewer) to duplicate dPCR measurements of Influenza A virus at the municipal treatment plant of Geneva, Switzerland (Oct 20 – Dec 26, 2022) using either 1) a normal likelihood (red), 2) a log-normal likelihood (orange), and 3) a dPCR-specific likelihood (blue) for the observations. Each panel shows the median (lines), and 50% and 95% credible intervals (strong and weakly shaded areas) of (A) posterior predictive distributions for dPCR measurements, (B) estimated viral loads in wastewater over time, and (C) the estimated effective reproduction number over time. Estimates shown beyond Dec 26, 2022 (vertical dotted line) are forecasted based on a continuation of the latest transmission dynamics. Panel A also shows the true observed (crosses) and future (stars) dPCR measurements.

**Fig S20.**
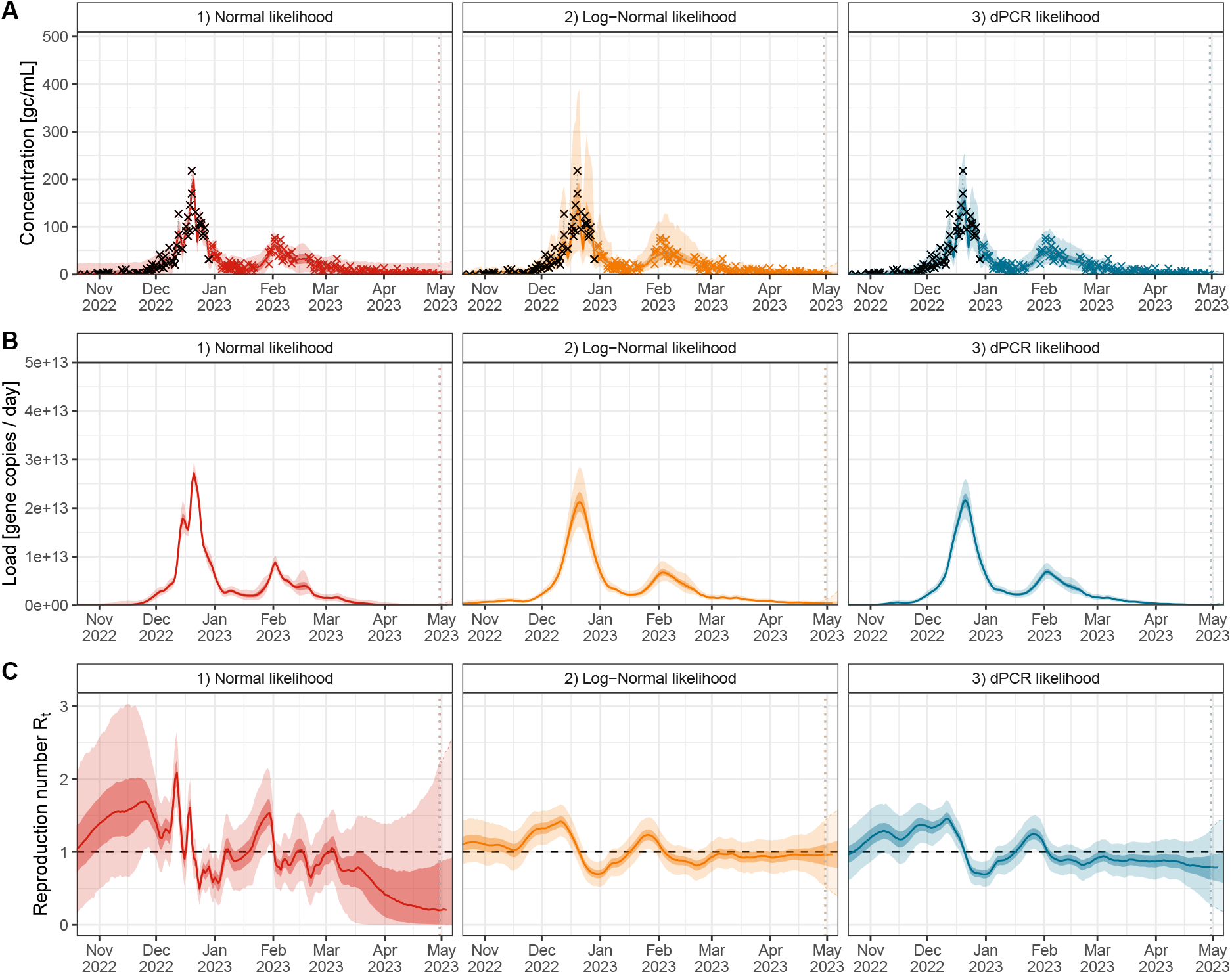
Comparison of likelihood functions on measurements of Influenza A virus concentrations at the municipal treatment plant of Geneva, Switzerland, Oct 20 – May 01, 2023: Shown are the results of fitting an epidemiological wastewater model (EpiSewer) to duplicate dPCR measurements of Influenza A virus at the municipal treatment plant of Geneva, Switzerland (Oct 20 – May 01, 2023) using either 1) a normal likelihood (red), 2) a log-normal likelihood (orange), and 3) a dPCR-specific likelihood (blue) for the observations. Each panel shows the median (lines), and 50% and 95% credible intervals (strong and weakly shaded areas) of (A) posterior predictive distributions for dPCR measurements, (B) estimated viral loads in wastewater over time, and (C) the estimated effective reproduction number over time. Estimates shown beyond Dec 26, 2022 (vertical dotted line) are forecasted based on a continuation of the latest transmission dynamics. Panel A also shows the true observed (crosses) and future (stars) dPCR measurements.

**Fig S21.**
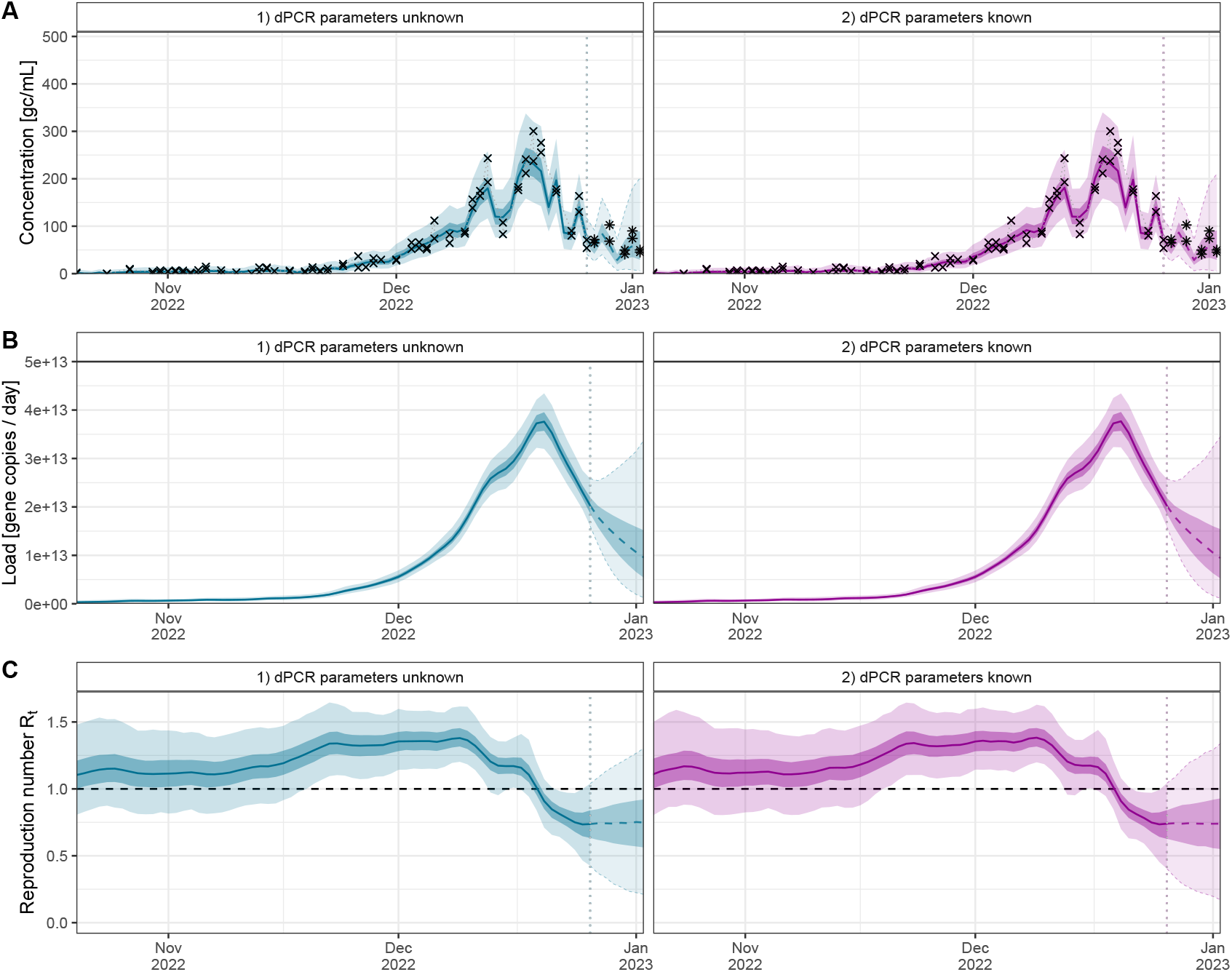
Comparison of inference and short-term forecasts from wastewater measurements using a dPCR-specific likelihood with and without knowledge of the dPCR parameters: Shown are the results of fitting an epidemiological wastewater model (EpiSewer) to duplicate dPCR measurements of Influenza A virus at the municipal treatment plant of Zurich, Switzerland (Oct 20 – Dec 26, 2022) using a dPCR-specific likelihood where 1) broad priors are used for the number of total partitions and the conversion factor (blue) and 2) the number of total partitions and the conversion factor are fixed to their true values (purple). Each panel shows the median (lines), and 50% and 95% credible intervals (strong and weakly shaded areas) of (A) posterior predictive distributions for dPCR measurements, (B) estimated viral loads in wastewater over time, and (C) the estimated effective reproduction number over time. Estimates shown beyond Dec 26, 2022 (vertical dotted line) are forecasted based on a continuation of the latest transmission dynamics. Panel A also shows the true observed (crosses) and future (stars) dPCR measurements.

